# Absence of GdX/UBL4A protects against inflammatory bowel diseases by regulating NF-κB signaling in DCs and macrophages

**DOI:** 10.1101/376103

**Authors:** Chunxiao Liu, Yifan Zhou, Mengdi Li, Ying Wang, Shigao Yang, Yarui Feng, Yinyin Wang, Yangmeng Wang, Fangli Ren, Jun Li, Zhongjun Dong, Y Eugene Chin, Xinyuan Fu, Li Wu, Zhijie Chang

**Author notes:** These authors contributed equally to this work (Chunxiao Liu, Yifan Zhou). Corresponding authors: Zhijie Chang; Li Wu;.

## Abstract

Nuclear factor-kappa B (NF-κB) activation is critical for innate immune responses. Here we report that the UBL4A (Ubiquitin-like protein 4A, also named GdX) enhances dendritic cells (DCs) and macrophages (Mφ)-mediated innate immune defenses by positively regulating NF-κB signaling. GdX-deficient mice were resistant to LPS-induced endotoxin shock and DSS-induced colitis. DC- or Mφ-specific GdX-deficient mice displayed alleviated mucosal inflammation, and the production of pro-inflammatory cytokines by GdX-deficient DCs and Mφ was reduced. Mechanistically, we found that PTPN2 (TC45) and PP2A form a complex with RelA (p65) to mediate its dephosphorylation whereas GdX interrupts the TC45/PP2A/p65 complex formation and restrict p65 dephosphorylation by trapping TC45. Our study provides a mechanism by which NF-κB signaling is positively regulated by an adaptor protein GdX in DC or Mφ to maintain the innate immune response. Targeting GdX could be a strategy to reduce over-activated immune response in inflammatory diseases.

## Introduction

NF-κB signaling is important in innate immune responses and is mediated by innate immune cells, including dendritic cells (DCs) and macrophages (Mφ). DCs and Mφ express pattern recognition receptors such as toll-like receptors (TLRs) to activate NF-κB, leading to the production of pro-inflammatory cytokines, including IL-1β, IL-6, and TNF-α during microorganism infection (Aderem and Ulevitch, 2000; Kawai and Akira, 2011). Dysregulation of NF-κB signaling is associated with a variety of human diseases, such as inflammatory bowel diseases (IBDs) (Molodecky et al., 2012).

The NF-κB signaling pathway is activated by TNF-α and TLR agonists (Hayden and Ghosh, 2008). In both situations, TRAF6 passes the baton to IKK for activation, leading to the phosphorylation and ubiquitination–proteasome dependent degradation of IκBα. Then, IκB-released p65 and p50 then form a dimer and translocate into the nucleus where targeted gene transcription is initiated (Ahmad et al., 2015). During the process, both IκB and p65 are phosphorylated by IKK. p65 phosphorylation at Ser276 and Ser536 by IKK or PKAc in the cytoplasm or by MSK1/2 or RSK-1 in the nucleus stabilizes their IκB free activation status (Oeckinghaus and Ghosh, 2009). Conversly, NF-κB signaling must be downregulated after its activation. One example by which NF-κB signaling is shut down during inflammation is through microorganisms that may trigger protein tyrosine phosphatases (PTPs) in immune cells (Pierce et al., 2011).

PTPs trigger p65 dephosphorylation to terminate its transcriptional activity (Ghosh and Hayden, 2008). Serine/threonine protein phosphatase 1 (PP1), PP2A, PP4 and WIP1 (Chew et al., 2009; Li et al., 2008; Li et al., 2006; Yang et al., 2001) have been identified to dephosphorylate p65. In particular, PP2A was identified as a specific regulator of NF-κB signaling by either suppressing the NF-κB transcriptional activity or reducing its binding ability to DNA. Dephosphorylation of NF-κB by PP2A leads to inhibition of NF-κB transcriptional activity, involved in chemokines or cytokines induction in astrocytes (Li et al., 2006). Recently, natural compounds isoliensine (Shu et al., 2016), rographolide (Hsieh et al., 2011), and the hydrophilic alpha-tocopherol derivative, PMC (Hsieh et al., 2014), are found to inhibit NF-κB activity by regulating its dephosphorylation through activation of PP2A. Overall, manipulation of PP2A activity regulates NF-κB activity on-and-off regulation in immune cells.

GdX is an X-linked gene in the G6PD cluster at Xq28; also named UBL4A (Ubiquitin-like protein 4A) that encodes a small protein with an N-terminal ubiquitin-like domain (Toniolo et al., 1988; Wang et al., 2014). GdX has also been reported to regulate ER stress responses for protein folding (Xu et al., 2012; Xu et al., 2013) and also to promote Akt activation through interacting with Arp2/3 in the cytoplasm (Zhao et al., 2015). While studying the regulation of transcription factor STAT3, we discovered GdX-mediated tyrosine phosphatase TC45 recruitment for STAT3 dephosphorylation (Wang et al., 2014). Surprisingly, in this study, we found that GdX-deficient mice (Wang et al., 2012) were resistant to LPS-induced endotoxin shock and DSS-induced colitis, suggesting a critical effect of GdX in NF-κB signaling pathway modulation. Here, we describe the mechanism of GdX in the regulation of p65 dephosphorylation and maintenance of p65 in a hyper-active status during inflammatory responses.

## Results

### GdX positively regulates the innate immune responses

To investigate the role of GdX in innate immune responses, a lethal dosage of LPS (30 mg/kg) was injected intraperitoneally to GdX^+/Y^ and GdX^−/Y^ mice (Wang et al., 2012). A significantly higher survival rate was observed in GdX^−/Y^ mice than in the control mice (GdX^+/Y^) (Figure 1A). We further compared the gene expression profiles of splenocytes from GdX^+/Y^ and GdX^−/Y^ mice upon LPS stimulation. Interestingly, RNA-sequencing analysis indicated a number of NF-κB targeting genes, such as IL-6, IL-1a and IL-12, have dramatic higher expressions in the splenocytes of GdX^+/Y^ mice than that of GdX^−/Y^ mice (Figure 1B, Figure 1-figure supplement 1A). Moreover, gene ontology (GO) analysis of GdX function in LPS-treated mice showed one of the most significantly enriched biological process is related to inflammatory response (Figure 1-figure supplement 1B). These results were further validated by testing the serum levels of IL-6 (Figure 1C) and TNF-α (Figure 1D), which are two important NF-κB targeted genes involved in innate immune responses. Correspondingly, lower mRNA levels of IL-6, TNF-α and IL-1β were detected in the splenocytes from GdX^−/Y^ mice compared to those from the control mice (Figure 1-figure supplement 1C-F), and was also validated in the thymus (Figure 1-figure supplement 1G) and liver (Figure 1-figure supplement 1H). These results suggest that GdX^−/Y^ mice were resistant to LPS challenge, implying a positive role for GdX in regulating the innate immune responses.

**Figure 1.**
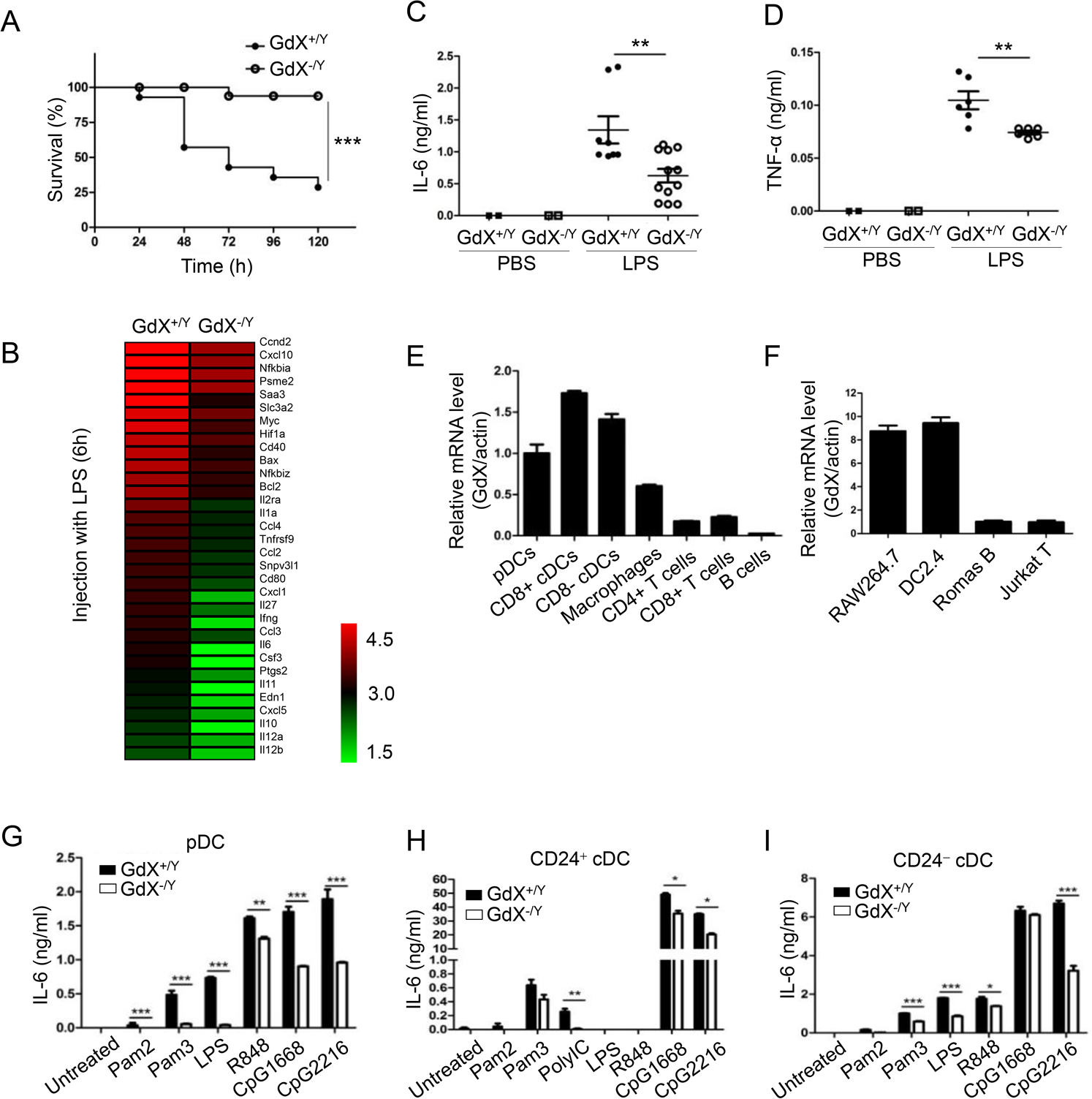
GdX deletion attenuates LPS-induced inflammatory responses *in vivo*. (A) GdX-deficient mice (GdX^−/Y^) were resistant to LPS challenge. Survival rates of GdX^−/Y^ and their wildtype (GdX^+/Y^) littermates (n=15) were observed after challenged with a lethal dosage of LPS (30 mg/kg body weight, ip). (B) A number of NF-κB target genes had significantly lower expressions in GdX^−/Y^ mice than control mice after LPS injection. The heat map of downregulated NF-κB target genes (list of 32 genes, absolute log2 >1, false discovery rate <0.05, p<0.05) in GdX^−/Y^mice (n=2) and wildtype littermates (n=2) injected with LPS. The mice were injected with LPS (20 mg/kg body weight, ip), and 6 hrs after injection, RNA was extracted from their splenocytes and performed the RNA-Seq analysis. (C-D) GdX deletion decreased the serum concentrations of pro-inflammatory cytokines under acute inflammation. Levels of IL-6 and TNF-α in serum from GdX^+/Y^ and GdX^−/Y^ mice (n≥6) were examined by ELISA after LPS challenge (20 mg/kg body weight; ip). (E) GdX is abundantly expressed in myeloid cells. mRNA levels of GdX were measured by real-time PCR in different immune cells. (F) GdX is highly expressed in RAW264.7 and DC2.4 cell lines. (E-F) The values were the mean ± SEM of three technical repeats. Data showed one representative experiment of three performed. (G-I) Different subtypes of FLDCs from GdX^+/Y^ and GdX^−/Y^ mice were used to measure the productions of IL-6 in response to indicated TLR ligand stimulation. The values were the mean of two biological repeats. Data showed the representative one of three repeats. *, p < 0.05; **, p < 0.01; ***, p < 0.001.

To address whether the resistance of GdX^−/Y^ mice was due to a change of immune cell homeostasis, we compared the immune cell profiles of GdX^−/Y^ mice and their control littermates. The results showed that the percentage of DCs, Mφs, B cells and T cells were similar in the lymphoid organs of GdX^+/Y^ and GdX^−/Y^ mice (Figure 1-figure supplement 1I). These results indicate that GdX is dispensable for immune cell development, suggesting that GdX might play a role in regulating the function of the immune cells.

### GdX promotes the production of inflammatory cytokines by DCs and Mφ

To reveal the role of GdX in immune cells, we compared the mRNA level of GdX in different immune cell types. The results showed that GdX was expressed abundantly in DCs (pDCs and cDCs) from the spleen, Mφ from BM, but weakly in T cells (CD4^+^ and CD8^+^), and rarely in B cells from the spleen (Figure 1E). Consistently, GdX was highly expressed in RAW264.7 and DC2.4 cell lines, but low in Romas B and Jurkat T cells (Figure 1F). These results suggest that GdX might play a role in myeloid cells, in particular, DCs and Mφ.

We then investigated whether GdX deficiency affected inflammatory cytokine production by DCs and Mφ. The Fms-like tyrosine kinase 3 ligand (Flt3L) supplemented BM cultured DCs (FLDCs, including pDCs, CD24^+^ cDCs and CD24^−^ cDCs, equivalent to splenic CD8α^+^ cDCs and CD8α^−^ cDCs respectively in the steady state) were stimulated with a panel of TLR ligands *in vitro* and examined for IL-6, IL-12, and other cytokines and chemokines. The results showed that the pDCs, CD24^+^ cDCs and CD24^−^ cDCs from GdX-deficient mice produced considerably lower amounts of IL-6 (Figure 1G-I) and IL-12 (Figure 1-figure supplement 1J-L) than that from WT mice in response to different stimulations. Simultaneously, the production of IFN-λ was dramatically decreased in pDCs from GdX-deficient mice in response to CpG2216 (Figure 1-figure supplement 1M). In addition, GdX deficiency slightly impaired the production of MIP-1α and RANTES by CD24^−^ cDCs (Figure 1-figure supplement 1N, O). In line with these findings, deletion of GdX significantly decreased LPS-induced secretion of IL-6 and TNF-α by GMDCs (Granulocyte-macrophage colony-stimulating factor-induced DCs) (Figure 1-figure supplement 1P, Q) and BMDMs (bone marrow-derived Mφ) (Figure 1-figure supplement 1R, S). These results suggest that GdX deficiency attenuates the production of inflammatory cytokines by myeloid cells.

Additionally, we confirmed that deletion of GdX had no significant effect on the cell viability (Figure 1-figure supplement 1T), expression of TLR2/4/9 (data not shown), or antigen presentation ability of the myeloid cells (data not shown). We speculated that the reduced inflammatory cytokine production by DCs and Mφ in GdX-deficient mice might be due to the regulation of TLR signaling.

### GdX facilitates the NF-κB signaling by promoting p65 activation

The NF-κB signaling pathway is considered a protypical pro-inflammatory pathway downstream of TLR activation. The regulatory role of GdX on the NF-κB signaling was examined by transfecting the NF-κB luciferase reporter (κB-luc) along with MyD88, TRAF6, IKKβ or p65, key components in the NF-κB signaling pathway, in HEK293T cells. The results showed that the luciferase activities were further increased by over-expression of GdX when MyD88 (Figure 2A), TRAF6 (Figure 2B), IKKβ (Figure 2C) or p65 (Figure 2D) was co-transfected. Over-expression of GdX also promoted NF-κB activation by TNF-α (Figure 2-figure supplement 1A) or LPS/TLR4 (Figure 2-figure supplement 1B), but inhibited STAT3 activity as we reported previously (Figure 2-figure supplement 1C) (Wang et al., 2014). In contrast, depletion of GdX using siRNAs inhibited NF-κB activation (Figure 2-figure supplement 1D-H). These results strongly suggest that GdX directly regulates p65 rather than the upstream components of the NF-κB signaling pathway.

**Figure 2.**
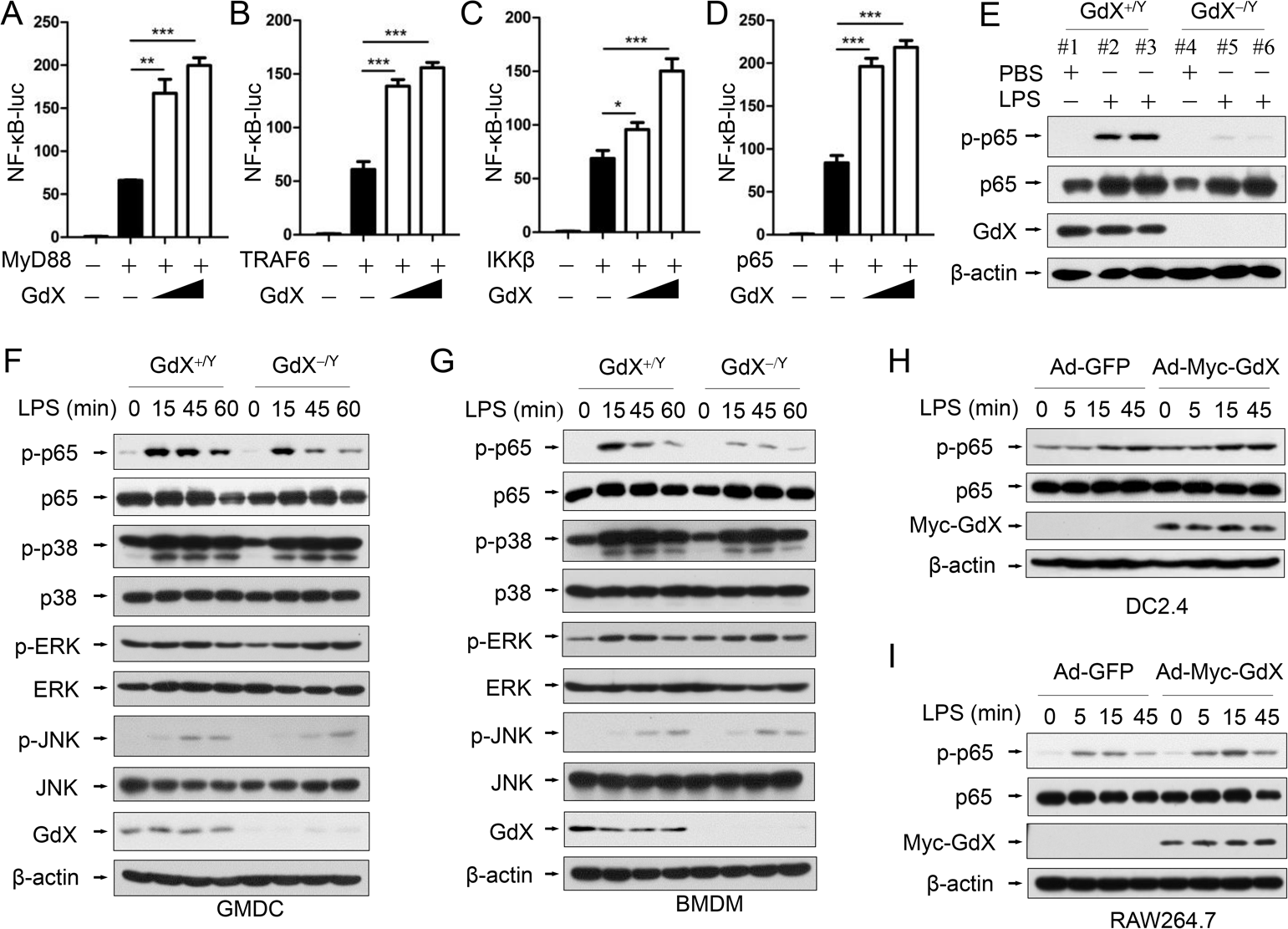
GdX positively regulates NF-κB signaling by targeting p65. (A-D) GdX promoted the transcriptional activity of NF-κB. HEK293T cells were transfected with NF-κB response luciferase reporter (NF-κB-luc), together with MyD88 (A), TRAF6 (B), IKKβ (C), or p65 (D), along with different dosages of GdX. Luciferase activity was measured at 36 h after transfection and the results were presented as mean ± SEM from three repeats. *, p < 0.05; **, p < 0.01; ***, p < 0.001. (E) The level of p-p65 was significantly lower in splenocytes of GdX^−/Y^ mice than that of GdX^+/Y^ mice. Individual mouse was labeled with *#* and number. (F and G) Deletion of GdX resulted in decreased phosphorylation of p65 in response to LPS. Levels of indicated proteins in GMDCs (F) and BMDMs (G) after LPS treatments were examined by WB. (H and I) Over-expression of GdX increased the levels of p-p65. DC2.4 (H) or RAW264.7 (I) cells infected with an adenovirus expressing GFP (Ad-GFP) or Myc-GdX (Ad-Myc-GdX) were stimulated with LPS (100 ng/mL). All of the in vitro experiments were repeated three times, and the results shown were representative.

To investigate the effect of GdX on p65, the splenocytes from GdX^−/Y^ mice, challenged with LPS for 16 h, were collected for western blot analyses. The results showed that p65 serine phosphorylation was dramatically decreased in the splenocytes obtained from GdX^−/Y^ mice, compared with that in GdX^+/Y^ mice (Figure 2E). We then treated the GMDCs and BMDMs of GdX^+/Y^ (WT) or GdX^−/Y^ mice with LPS, and examined the activation of different signaling pathways involved in TLR activation. The p65 phosphorylation intensity was dramatically decreased in both GMDCs (Figure 2F, top panel) and BMDMs (Figure 2G, top panel) of GdX^−/Y^ mice. However, levels of phospho-p38, phospho-ERK and phospho-JNK are comparable (Figure 2F, 2G, middle panels). These results suggest that GdX specifically regulates the phosphorylation of p65 in DCs and Mφ. Additionally, p65 phosphorylation induced by LPS was increased in GdX over-expressed DC2.4 and RAW264.7 cells (Figure 2H, 2I). These results suggest that GdX participates in TLR-activated NF-κB pathway, up-regulating the phosphorylation of p65 in both *in vitro* and *in vivo* models, promoting the pro-inflammatory cytokine production by DCs and Mφ.

### GdX disrupts the TC45/p65 complex formation

Since GdX had no direct interaction with p65 (Figure 3-figure supplement 1A), we hypothesized that GdX regulates the phosphorylation status of p65 by controlling a protein kinase or phosphatase. We focused on IKKα, IKKβ, IKKε, canonical and non-canonical kinases known to be important for the phosphorylation of p65 (Hoesel and Schmid, 2013; Sugimoto et al., 2008); PP2A, a serine phosphatase that dephosphorylates p-p65 (phosphorylated p65) (London et al., 2010; Nathan, 2002); WIP1, a PP2C family number (Chew et al., 2009), and TC45, a tyrosine phosphatase that interacts with GdX (Wang et al., 2014). Immunoprecipitation (IP) experiments demonstrated that GdX interacted only with TC45 among these kinases and phosphatases (Figure 3A), suggesting TC45 might associate with the regulation of p65 phosphorylation by GdX.

**Figure 3.**
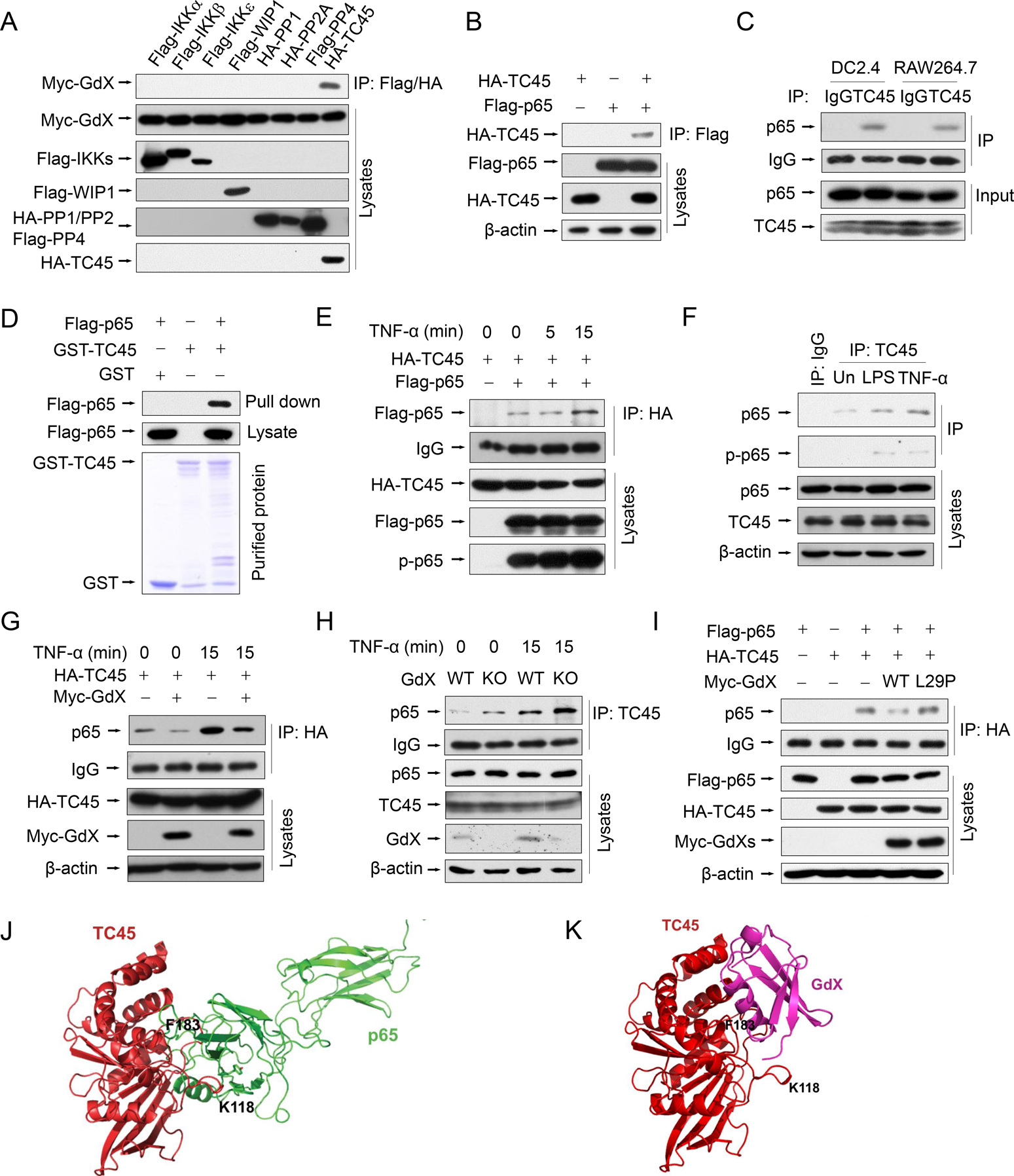
GdX blocks the interaction of TC45 with p65. (A) GdX specific interacted with TC45. Myc-GdX was co-expressed with Flag-tagged IKKa, IKKβ, IKKε, WIP1, PP4 or HA-tagged PP1, PP2A and TC45 in HEK293T cells. Immunoprecipitation (IP) experiments were performed using an anti-Flag or HA antibody. (B) HA-TC45 interacted with Flag-p65. HEK293T cells were transfected for IP with an anti-Flag antibody. (C) The association of endogenous TC45 and p65 was detected in immune cell lines. Lysates from DC2.4 or RAW264.7 cells were subjected to IP experiments with an antibody against TC45. (D) Flag-p65 interacted with GST-TC45 purified from *E. coli*. (E) The interaction of Flag-p65 and HA-TC45 was increased under TNF-α stimulation (10 ng/mL) for 15 min. (F) The interaction of endogenous TC45 and p65 was increased under LPS or TNF-α stimulation. IP experiments were performed by an anti-TC45 antibody. (G) Over-expression of GdX disrupted the association of TC45 and p65. HEK293T cells were used for IP experiments after transfected with HA-TC45 in the presence or absence of Myc-GdX. (H) The association of endogenous TC45 and p65 was increased in GdX-deficient (KO) splenocytes comparing with that in the wildtype (WT) cells. (I) GdX(L29P) mutant failed to block the interaction of p65 and TC45. (J-K) A molecular docking analysis shows the interaction surface of TC45 with the N-terminus of p65 (J) and GdX (K). F183 in TC45 maintains a core residue for the interaction with p65 and GdX although GdX shifts slightly to one side in the interaction (K). All of the in vitro experiments were repeated three times, and the results shown were representative.

We then questioned whether TC45 could directly interact with p65. Co-IP analyses revealed that Flag-tagged p65 was able to precipitate with HA-tagged TC45 (Figure 3B), and reciprocally, HA-TC45 pulled down Flag-tagged p65 (Figure 3-figure supplement 1B). Consistently, the endogenous TC45 protein associated with p65 in both DC2.4 and RAW264.7 cells (Figure 3C). In addition, purified proteins GST-TC45 pulled down Flag-p65, suggesting a direct interaction of TC45 with p65 (Figure 3D). Furthermore, the interaction of p65 and TC45 was enhanced after TNF-α stimulation in both the over-expression (Figure 3E) and endogenous conditions (Figure 3F). Interestingly, when Myc-GdX, HA-TC45 and Flag-p65 were co-expressed together, we observed that HA-TC45 precipitated down both Myc-GdX and Flag-p65, but Myc-GdX failed to precipitate down Flag-p65 (although Myc-GdX precipitated down HA-TC45), and Flag-p65 failed to precipitate down Myc-GdX (Figure 3-figure supplement 1C). These results indicated that TC45 exclusively forms a complex with p65 or GdX.

To analyze whether GdX affects the association of TC45 and p65, IP experiments were performed using HA-TC45 and endogenous p65 protein in the absence or presence of Myc-GdX in 293T cells. The results showed that the interaction of p65 and HA-TC45 was significantly decreased in the presence of Myc-GdX in conditions with or without TNF-α stimulation (Figure 3G). Consistently, we observed that the interaction of endogenous TC45 with p65 was dramatically increased in the splenocytes from GdX^−/Y^ mice (Figure 3H). Similar results were obtained in GMDCs (Figure 3-figure supplement 1D) and BMDMs (Figure 3-figure supplement 1E) and suggest that GdX inhibits TC45 binding to p65.

To reveal whether GdX-impaired interaction of TC45 and p65 is due to the interaction of GdX with TC45, we recruited a mutant GdX, GdX(L29P), which lacks the ability to interact with TC45 (Wang et al., 2014). IP experiments demonstrated that GdX(L29P) failed to disrupt the interaction of TC45 and p65 (Figure 3I). Therefore, we conclude that GdX interacts with TC45 and then disrupts the interaction of TC45 with p65.

Furthermore, we performed a molecular docking analysis to show the detailed interaction of TC45 with p65 (Figure 3J) and GdX (Figure 3K). The results showed that the surface of TC45 for the interaction with GdX is the same as it interacts with p65 although GdX interacts on a slight shift site. In particular, residue F183 in TC45 faces to the interacting sites of p65 and GdX (Figure 3J and 3K). Therefore, the interaction of GdX with TC45 occupies F183 and disrupts the interaction of p65 with TC45 (Figure 3-figure supplement 1F). This structure base of interaction explains the mechanism for the exclusive interaction of TC45 with GdX or p65.

### GdX prolongs p65 phosphorylation

Since GdX inhibits TC45 binding to p65, we speculated that the altered p-p65 level by GdX might be due to an altered dephosphorylation process. To test this possibility, we stimulated the GMDCs and BMDMs with LPS, and then we withdrew LPS and allowed cells to undergo starvation to induce protein dephosphorylation. Western blot analyses demonstrated that the level of p-p65 was slightly decreased in GMDCs and BMDMs from WT mice but quickly decreased in the cells from GdX^−/Y^ mice when the cells were subject to starvation for different times (Figure 4A, 4B). Similar results were observed in the splenocytes from WT and GdX^−/Y^ mice (Figure 4-figure supplement 1A). Reciprocally, when GdX was over-expressed by an adenovirus in DC2.4 and Raw264.7 cells, which were challenged by LPS for 30 min, high level of p-p65 remained for 60 min after starvation (Figure 4C, 4D). Over-expression of GdX also increased the level of p-p65 after TNF-α stimulation in 293T cells (Figure 4-figure supplement 1B). These results demonstrated that deletion of GdX shortened, while over-expression of GdX extended, the maintaining time of p-p65. In consistency with the prolonged p-p65 levels, we further observed that p65 occupied the promoter of IL-6 for a longer time (Figure 4-figure supplement 1C, D) and remained in the nucleus after starvation for 120 min when GdX was over-expressed, while it redistributed into the cytoplasm after starvation for 60 min in the control cells (Figure E). These results suggested that GdX abrogates the dephosphorylation process of p-p65.

**Figure 4.**
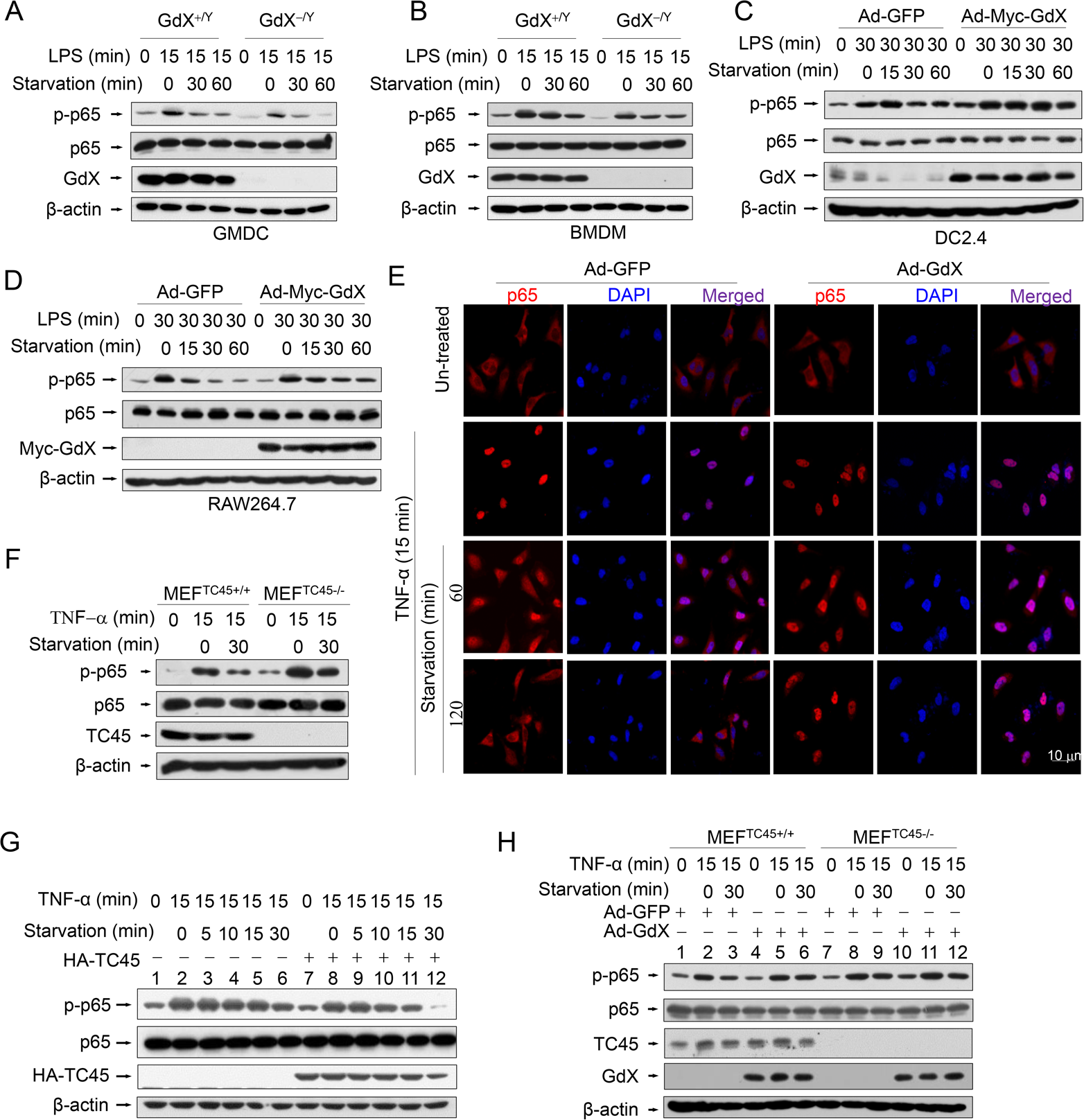
GdX maintains the phosphorylation of p65 by counteracting TC45. (A and B) Depletion of GdX led to increased dephosphorylation of p65 in GMDC (A) and BMDM (B) cells. Cells derived from GdX^+/Y^ or GdX^−/Y^ mice were treated with LPS (100ng/mL) and then subjected to starvation at different times for dephosphorylation. (C and D) GdX sustained the level of p-p65 in DC2.4 (C) and RAW264.7 (D) cell lines. Cells were treated with LPS (100ng/mL) after infection with an adenovirus expressing GFP or GdX and then subjected to starvation for indicated times. (E) HeLa cells were infected with either Ad–GFP (an adenovirus expressing GFP) or Ad–GdX (an adenovirus expressing GdX) for 36 h, and then treated with TNF-α for indicated times. Subcellular localization of p65 was analyzed by immunofluorescence confocal microscopy with a rabbit anti-p65 antibody (red). DAPI was used to stain the nucleus. Scale bar, 10 μm. (F) Dephosphorylation of p-p65 was inhibited in TC45-depleted cells. p-p65 levels were examined in MEF^TC45+/+^ and MEF^TC45−/−^ cells treated with or without TNF-α (10 ng/mL) for 15 min, followed by starvation for 30 min before harvesting. (G) TC45 accelerated p65 dephosphorylation. p-p65 levels were examined in HEK293T cells, which were treated with TNF-α for 15 min and then subjected to starvation, in the presence or absence of HA-TC45. (H) GdX failed to maintain the p-p65 level in TC45-deficient cells. All of the in vitro experiments were repeated three times, and the results shown were representative.

Given that TC45 interacts with p65, we speculated that TC45 might mediate the dephosphorylation of p65. Indeed, deletion of TC45 in MEFs showed a significantly prolonged phosphorylation of p65 (Figure 4F) while over-expression of TC45 dramatically decreased the level of p-p65 after starvation for 30 min (Figure 4G, lanes 12 and 6). Since GdX interrupts the interaction of TC45 with p65, we questioned whether GdX affects the dephosphorylation of p65 via blocking TC45. For this purpose, we used MEFs with a TC45 deletion, under the over-expression of GdX. We observed that, over-expression of GdX maintained the level of p-p65 after starvation for 30 min in the WT cells (Figure 4H, comparing lane 6 to lane 3), confirming that GdX impairs the dephosphorylation of p-p65. However, over-expression of GdX failed to further increase the level of p-p65 under the starvation condition when TC45 was deleted (Figure 4H, comparing lanes 12 to 9), indicating that GdX is unable to regulate the p-p65 level without TC45. These results suggest that GdX regulates the dephosphorylation of p65 via TC45. Taken together, we conclude that GdX-elevated phosphorylation of p65 is due to the interruption of TC45 from binding to p65.

### Residue Y100 in p65 is critical for TC45 to mediate p65 dephosphorylation

We next mapped the region for the interaction of p65 with TC45. IP experiments (Figure 5-figure supplement 1A) showed that truncated p65-n6 (where 1–90 amino acids remained) failed to interact with TC45, whereas other truncated forms of p65 maintained strong interactions with TC45 (Figure 5-figure supplement 1B). These data suggested that the region from amino acid 90 to 170 is essential for p65 to interact with TC45.

Since TC45 is a tyrosine phosphatase, we checked the conserved residues in this region and identified two tyrosine residues Y100 and Y152 (Figure 5A). We speculated that these two tyrosine residues might be critical for TC45-mediated p65 dephosphorylation. To examine this hypothesis, we mutated these two residues into phenylalanine (F). Luciferase experiments indicated that p65(Y100F) had a decreased activity on the NF-κB reporter whereas p65(Y152F) remained the same activity as wild type p65 (Figure 5B, black columns). Interestingly, over-expression of TC45 failed to inhibit p65(Y100F)-induced luciferase activity but remained to inhibit p65(Y152F)-induced activity (Figure 5B). These results suggest that TC45 inhibits the activity of p65 through Y100. Simultaneously, we observed that over-expression of GdX elevated the reporter activity mediated by p65 and p65(Y152F) but had no effect on p65(Y100F)-induced activation (Figure 5C). Consistent with these observations, over-expression of TC45 appeared to have no effect on the phosphorylation level of p65(Y100F), which though appeared lower than that of p65 (WT) and p65(Y152F) (Figure 5D). We further deciphered that p65(Y100F) lost the interaction with TC45 but retained the interaction with p65(Y152F) (Figure 5E). A molecular structure docking analysis suggests that this Y100 forms a link with K118 at TC45 to maintain a surface for the interaction of p65 with TC45 (Figure 5-figure supplement 1C, D). These results indicate that Y100 is a key residue for the interaction of p65 with TC45 and thereafter the regulation of p65 dephosphorylation by TC45.

**Figure 5.**
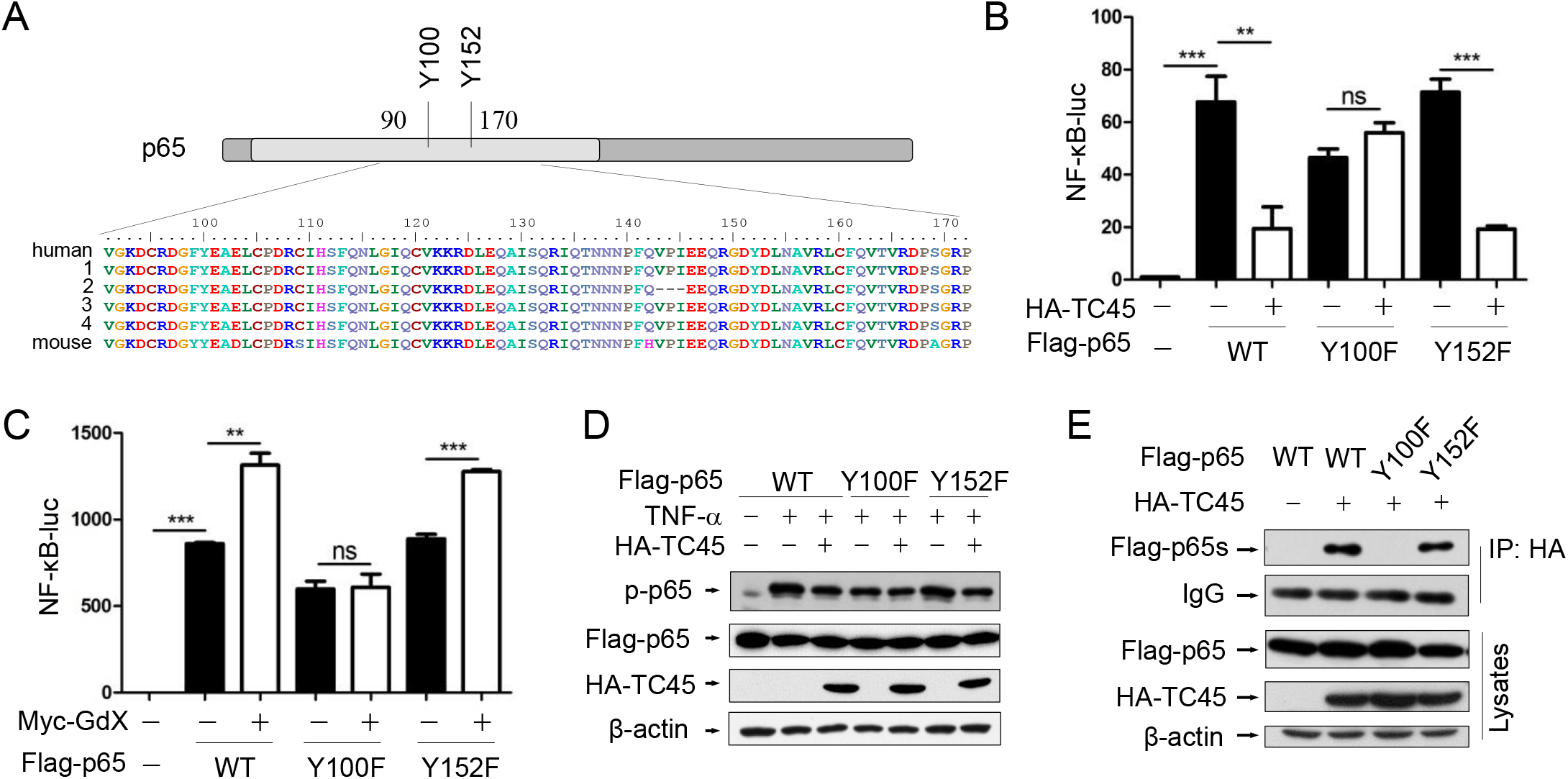
Tyrosine-100 in p65 is critical for TC45 function. (A) A schematic diagram showing two tyrosine (Y) residues in p65 in the region of amino acids 90-170. Both human and mouse sequences are showed. Numbers on the left indicated variants of p65 in human. (B) Y100 in p65 is critical for TC45 to suppress NF-κB signaling. Y: tyrosine; F: phenylalanine. (C) Y100 in p65 is critical for GdX to promote the activation of NF-κB signaling. Results were presented as mean ± SEM from three independent repeats. ns: no significant difference; *, p < 0.05; **, p < 0.01; ***, p < 0.001. (D) TC45 failed to promote the dephosphorylation of p65(Y100F) mutant. HEK-293T cells were transfected with the Flag-tagged p65, p65(Y100F), or p65(Y152F), in the presence or absence of HA-TC45. (E) TC45 failed to interact with p65(Y100F). HEK-293T cells were transfected with indicated plasmids. IP experiments were performed by using an anti-HA antibody. All of the in vitro experiments were repeated three times, and the results shown were representative. The results were presented as mean ± SEM. **, p < 0.01; ***, p < 0.001.

### TC45 recruits PP2A to dephosphorylate p65 at S536

To determine whether TC45 recruits a serine phosphatase to reduce p65 S536 phosphorylation, we screened several p65 S536 phosphatases. The results showed that over-expression of TC45 further decreased the phosphorylation of p65 based on PP2A, but not on other phosphatases including PP1, PP4 and WIP1 (Figure 6A), which implied that TC45 might mediate the dephosphorylation of p65 through PP2A. PP2A is the main source of phosphatase activity in the cell, which has been reported to interact with p65 (Yang et al., 2001). We then questioned whether TC45 affects the interaction of PP2A with p65. Intriguingly, we observed that TC45 enhanced the interaction of PP2A with p65 (Figure 6B). On the other hand, the interaction of PP2A with p65 was dramatically impaired when TC45 was knocked out (Figure 6C) or depleted by siRNA (Figure 6-figure supplement 1A). Moreover, we observed that over-expression of PP2A failed to mediate the dephosphorylation of p65 in TC45-deficient MEFs (Figure 6D, lanes 11-12), but strongly decreased the phosphorylation level in the WT cells (Figure 6D, lanes 5-6). These results suggest that TC45 is required for PP2A-mediated dephosphorylation of p-p65.

**Figure 6.**
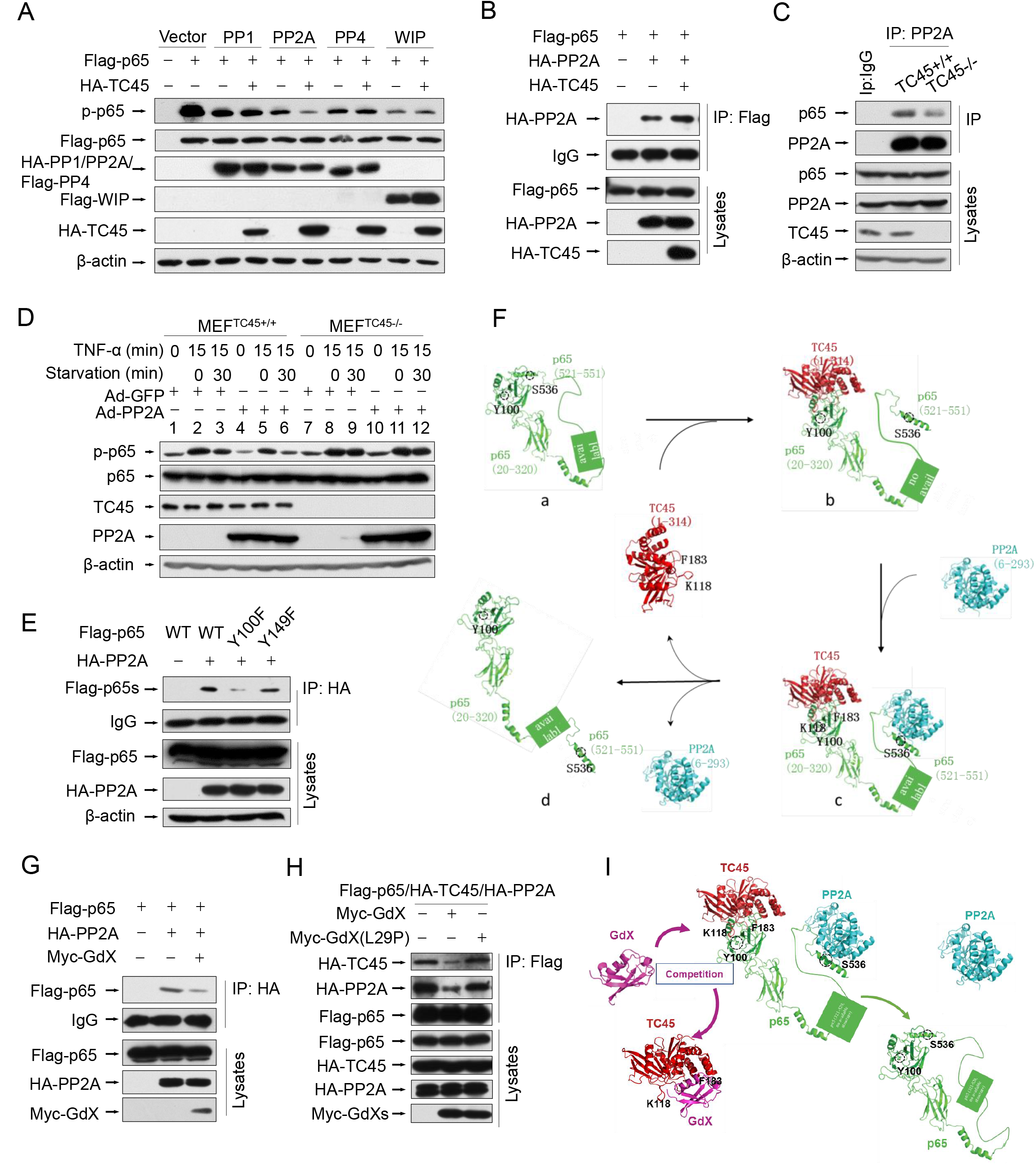
TC45 mediates PP2A to dephosphorylation p-p65, which is interrupted by GdX. (A) TC45 and PP2A synergistically dephosphorylate p-p65. HEK293T cells were co-transfected with different phosphatases and Flag-tagged p65 plasmids. (B) Overexpression of TC45 enhanced the interaction of PP2A and p65. (C) The endogenous interaction of PP2A and p65 was decreased in TC45-deficient cells. (D) PP2A failed to dephosphorylate p-p65 without TC45. p-p65 levels were examined in MEF^TC45+/+^ and MEF^TC45−/−^ cells infected with Ad-GdX or Ad-PP2A and treated with TNF-α (10 ng/mL) for 15 min, followed by TNF-α withdrawal for 30 min before harvesting. (E) Y100 is critical for the interaction of PP2A and p65. IP assays were performed after HEK-293T cells were transfected with the Flag-tagged p65, p65(Y100F), or p65(Y152F), as well as HA-PP2A for 24 h. (F) A molecular docking model showed TC45/p65/PP2A complex formation and disassociation. (a) The C-terminus of p65 is hided at the N-terminus of p65 to maintain an active form of p65. Without association of TC45 with p65 at its N-terminus, p65 remains a coated confirmation where the C-terminus, in particular S538, is hided. (b) TC45 starts to bind to the C-terminus of p65 and release the N-terminus of p65 from hiding. (c) PP2A gets a chance to associate with the released C-terminus and the heter-trimer complex of TC45/p65/PP2A is formed. In this complex, PP2A dephosphorylates S536 and finally maintains p65 unphosphorylated status (d). (G) GdX inhibited the interaction of p65 and PP2A. (H) GdX(L29P) mutant failed to decrease the interaction of PP2A and p65. HEK293T cells were transfected with the indicated plasmids for the IP experiment. (I) A model of the competition of GdX with TC45 to interact with p65. When TC45 interacts with the N-terminus of p65, the C-terminus of p65 is exposed to interact with PP2A. GdX interacts with TC45 and blocks its interaction with p65. In this way p65 maintains at its active form without interaction of PP2A. All of the in vitro experiments were repeated three times, and the results shown were representative.

Further results revealed that TC45, PP2A and p65 forms a hetero-trimer complex (Figure 6-figure supplement 1B). Interestingly, p65 (Y100F) showed much less interaction with PP2A (Figure E), suggesting that Y100 is critical for p65 to interact with both PP2A and TC45. Molecular docking analyses provided the structural base for the hetero-trimer complex, where TC45 binds to the N-terminus, and PP2A binds to the C-terminus separately (Figure 6Fc). In phosphoylated p65, the C-terminus is covered by the N-terminus (Figure 6Fa). When the C-terminus of p65 is released by TC45 (Figure 6Fb), PP2A associates with it and initiates the dephosphorylation of S536 (Figure 6Fc), leading to p65 dephosphorylation (Figure 6Fd).

Next, we determined whether GdX affects the interaction of PP2A with p65. Interestingly, IP experiments demonstrated that over-expression of Myc-GdX inhibited the interaction of HA-PP2A with Flag-p65 (Figure 6G). Consistently, we observed that the interaction of PP2A and p65 was dramatically increased when GdX was deleted in GMDCs (Figure 6-figure supplement 1C) and BMDMs (Figure 6-figure supplement 1D). To valid that the inhibitory role of GdX on the interaction of PP2A and p65 is due to its interaction with TC45, we used mutant GdX(L29P), which lost the ability to associate with TC45. The results showed that GdX(L29P) had a lesser influence on the interaction of PP2A and p65 (Figure 6-figure supplement 1E) and also the interaction of PP2A with TC45 (Figure 6-figure supplement 1F), similar to that as observed for its effect on the interaction of TC45 with p65 (Figure 3I). Consistent with the alteration of p65 phosphorylation (Figure 6A), we confirmed that TC45 further inhibited the NF-κB transcriptional activity based on over-expression of PP2A (Figure 6-figure supplement 1G). Furthermore, we observed that over-expression of Myc-GdX abrogated the complex of TC45/PP2A/p65 (Figure 6H). This rescued the p-p65 level, which was decreased by over-expression of HA-PP2A (Figure 6-figure supplement 1H), and was consistent with the results that GdX, but not GdX (L29P), rescued the NF-κB transcriptional activity inhibited by both TC45 (Figure 6-figure supplement 1I) and PP2A (Figure 6-figure supplement 1J). Based on the biochemical results, we proposed a model by molecular docking analysis. TC45 associates with p65 at the N-terminus via Y100, which leads to the release of the C-terminus of p65 to interact with PP2A. In this way, TC45, PP2A and p65 form a complex to dephosphorylate S536 of p65. When GdX sequestered TC45, the complex of TC45/PP2A/p65 is impaired and dephosphorylation of p65 is abrogated (Figure 6I).

### Specific deletion of GdX in DCs or Mφ protects the mice from colitis

The dysregulation of NF-κB signaling and innate immunity are closely associated with IBDs; thus we further examined the role of GdX in acute colitis. To evaluate the outcome of acute colitis in GdX^−/Y^ mice and their littermates, we used a model in which the mice were administrated with 3% DSS in drinking water for 6 days. GdX^−/Y^ mice exhibited a significantly decrease in body weight loss (Figure 7A), a longer colon length (Figure 7E, Figure 7-figure supplement 1A), and a markedly lower disease activity index (DAI) (Figure 7I) in comparison of control WT mice. The DAI is based on the magnitudes of body weight loss, diarrhea, and hemorrhage (Murano et al., 2000). Furthermore, histological analyses of the bowel tissue from the WT mice administered with DSS showed manifestations of inflammatory colitis, including loss of crypts, mucosal erosion, ulcers, and infiltration of inflammatory cells (Figure 7M, comparing b, c to a). In contrast, the specimens from the DSS-administered GdX^−/Y^ mice showed lower levels of inflammation, with marginal infiltration in the mucosa (Figure 7M, comparing e, f to d; also, comparing e to b and f to c).

**Figure 7.**
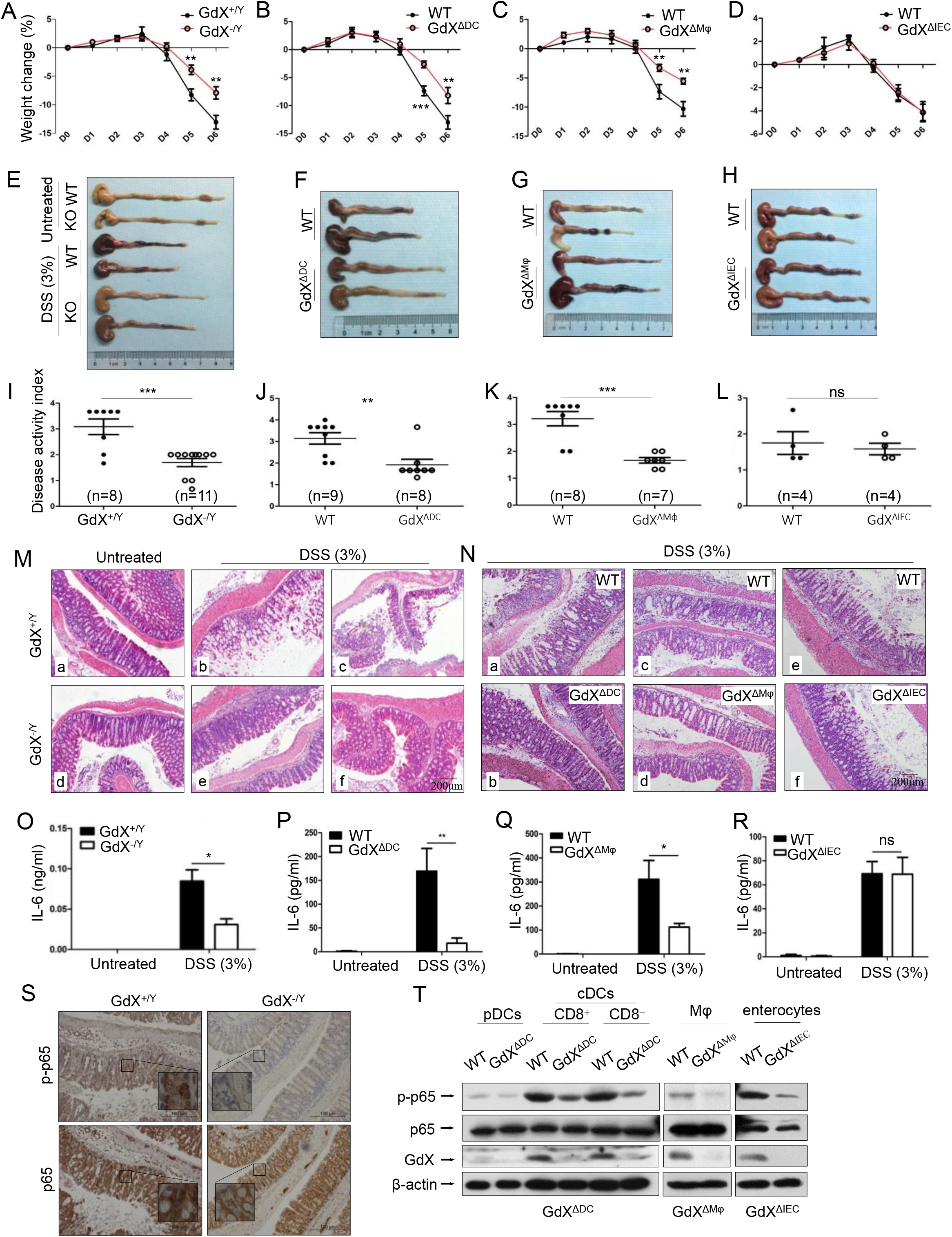
GdX-deficient mice show a reduced inflammatory colitis. Mice of GdX germline deletion (GdX^−/Y^), specific deletion in DC cells (GdX^ΔDC^), macrophages (GdX^ΔMφ^), epithelial cells (GdX^ΔIEC^) and their WT (Cre-negative) littermates were treated with 3% DSS in drinking water for 6 days. (A) Body weight of GdX^−/Y^ mice was decreased in comparison with that of GdX^+/Y^ mice at day 5 after DSS treatment. Body weight was measured daily and presented as means ± SEM. p value on day 5 was significant, p < 0.05 (n=6 for DSS-treated WT group, and n=12 for KO group). (B and C) Body weights of GdX^ΔDC^ (B) and GdX^ΔMφ^ (C) mice were decreased in comparison with that of WT mice after DSS treatment. n≥7 in each DSS-treated groups. (D) Body weight of GdX^ΔIEC^ mice was unchanged compared with that of WT mice after DSS treatment. n=4 in each DSS-treated groups. (E-H) The effect of GdX deletion on the colon lengths. The length of colon from GdX^−/Y^ (E), GdX^ΔDC^ (F), GdX^ΔMφ^ (G) and GdX^ΔIEC^ (H) mice and their WT (Cre-negative) littermates were determined on day 6 after DSS treatment. (I-L) The disease activity index (DAI) was determined on day 6. The DAI of GdX^−/Y^ (I), GdX^ΔDC^ (J), GdX^ΔMφ^ (K) and GdX^ΔIEC^ (L) mice were compared with their WT (Cre-negative) littermates. (M and N) Microscopic views of HE-stained colon sections. Tissue sections from GdX^−/Y^ (M), GdX^ΔDC^, GdX^ΔMφ^ and GdX^ΔIEC^ (N) mice and their WT (Cre-negative) littermates were subject to HE-staining. n≥4 in each DSS-treated groups. (O-R) The production of IL-6 was decreased in GdX^−/Y^, GdX^ΔDC^ and GdX^ΔMφ^ mice but not in GdX^ΔIEC^ mice. n≥4 in each DSS-treated groups. The serum IL-6 concentrations of the mice were compared with their WT (Cre-negative) littermates by ELISA. (S) Deletion of GdX resulted in decreased phosphorylation of p65 in IECs. Immuno-histochemical analyses of p-p65 (upper) and p65 (lower) in paraffin-embedded colon sections of GdX^+/Y^ (n=5) and GdX^−/Y^ mice (n=5) at day 6 after treatment with 3% DSS; nuclei were counterstained with hematoxylin (blue). Scale bars, 100 μm. (T) Examinations of p-p65 levels and GdX depletion efficiency in the spleen DCs of GdX^ΔDC^ mice and Mφ of GdX^ΔMφ^ mice. The DCs, Mφ were sorted by FACS and then protein levels were determined by WB. The data was the representative one of three repeats.

To further investigate the whether GdX regulate IBD development by innate immune cell-intrinsic mechanisms, we generated GdX conditional knockout mice in DCs (GdX^ΔDC^), Mφ (GdX^ΔMφ^) and intestine epithelial cells (IECs) (GdX^ΔIEC^) by crossing the CD11c-Cre, LysM-Cre or Villin-Cre mice to GdX^flox/flox^ mice. After administration with 3% DSS in drinking water for 6 days, GdX^ΔDC^, and GdX^ΔMφ^ mice lost less weight than littermate controls (Figure 7B, 7C), exhibited longer colons lengths (Figure 7F, 7G, Figure 7-figure supplement 1B, C), lower disease activity index (DAI) (Figure 7J, 7K), and significantly lower levels of inflammation observed via histological damage (Figure 7N, comparing b to a, d to c). The phenotypes of GdX^ΔDC^, and GdX^ΔMφ^ mice during acute colitis is similar with that observed in GdX^−/Y^ mice, suggesting deletion of GdX in DC and Mφ attenuate intestinal inflammation. However, GdX^ΔIEC^ mice did not show decreased intestinal inflammation (Figure 7D, H, L, N, e and f showed similar severe damages, Figure 7-figure supplement 1D), suggesting the deletion of GdX in IECs w,as not responsible for the decreased severity of colitis in GdX^−/Y^ mice. In addition, GdX^−/Y^ (Figure 7O), GdX^ΔDC^ (Figure 7P) and GdX^ΔMφ^ (Figure 7Q) mice showed significantly decreased levels of IL-6 in serum compared with the control mice, whereas GdX^ΔIEC^ mice maintained comparable levels of serum IL-6 to that of the control mice (Figure 7R) during DSS-induced colitis. These data suggest that the alleviated DSS-induced colitis in GdX^−/Y^ mice is largely due to functional changes of DCs and Mφ.

Inflammatory disorders in the gut are usually associated with disrupted homeostasis of T regulatory (T_reg_) and T helper 17 (Th17) cells, which can be induced by three types of myeloid cells including CD 11c^hi^CD11b^+^CD103^+^ DCs, CD11c^hi^CD11b^−^CD103+DCs and F4/80^+^CD11c^int^CD11b^+^CD103^−^ Mφ (Iwasaki, 2007). However, we did not find any changes in the frequencies of the intestinal T_reg_, Th17 cells and myeloid cell populations in GdX^ΔDC^ mice compared to WT mice after DSS treatment for 6 days (Figure 7-figure supplement 1E). These results supported the notion that the reduced severity of gut inflammation in GdX-deficient mice was largely due to the impaired production of inflammatory cytokine in the myeloid cells (DCs and Mφ), rather than the consequence of defects in myeloid cell development and Treg/Th17 cell balance.

As our aforementioned results suggested that GdX regulates dephosphorylation of p65, we examined the levels of p-p65 in the colons of the mice after DSS treatment. Immunohistochemical analyses demonstrated that p-p65 staining was much stronger in the nucleus in the colon section from WT mice than that from GdX^−/Y^ mice treated with DSS (Figure 7S, upper panels), while the non-phosphorylated p65 remained mainly in the cytoplasm of the cells from GdX^−/Y^ mice (Figure 7S, bottom panels). These results suggested that deletion of GdX impaired the activation of NF-κB in colon tissue. Consistently, we observed that specific deletion of GdX in DCs and Mφ significantly decreased the p-p65 levels during acute colitis (Figure 7T). Taken together, these data suggested that GdX deficiency alleviates the colon inflammation through regulation of NF-κB activity in DCs and Mφ.

## Discussion

In this study, we revealed a previously unrecognized regulatory mechanism of NF-κB signaling in innate immune cells. GdX forms a complex with tyrosine phosphatase TC45, traps PP2A by blocking its termination to warranty a sufficient activation of NF-κB (Figure 6I). GdX deletion impaired the production of pro-inflammatory cytokines by DCs and Mφ, protected mice against septic shock and acute colitis (Figure 8). Immune system requires a mechanism to fight against antigens rapidly and efficiently, where GdX functions as a ‘bodyguard’ in DCs and Mφ to maintain their sensitivity to pathogenic stimuli.

**Figure 8.**
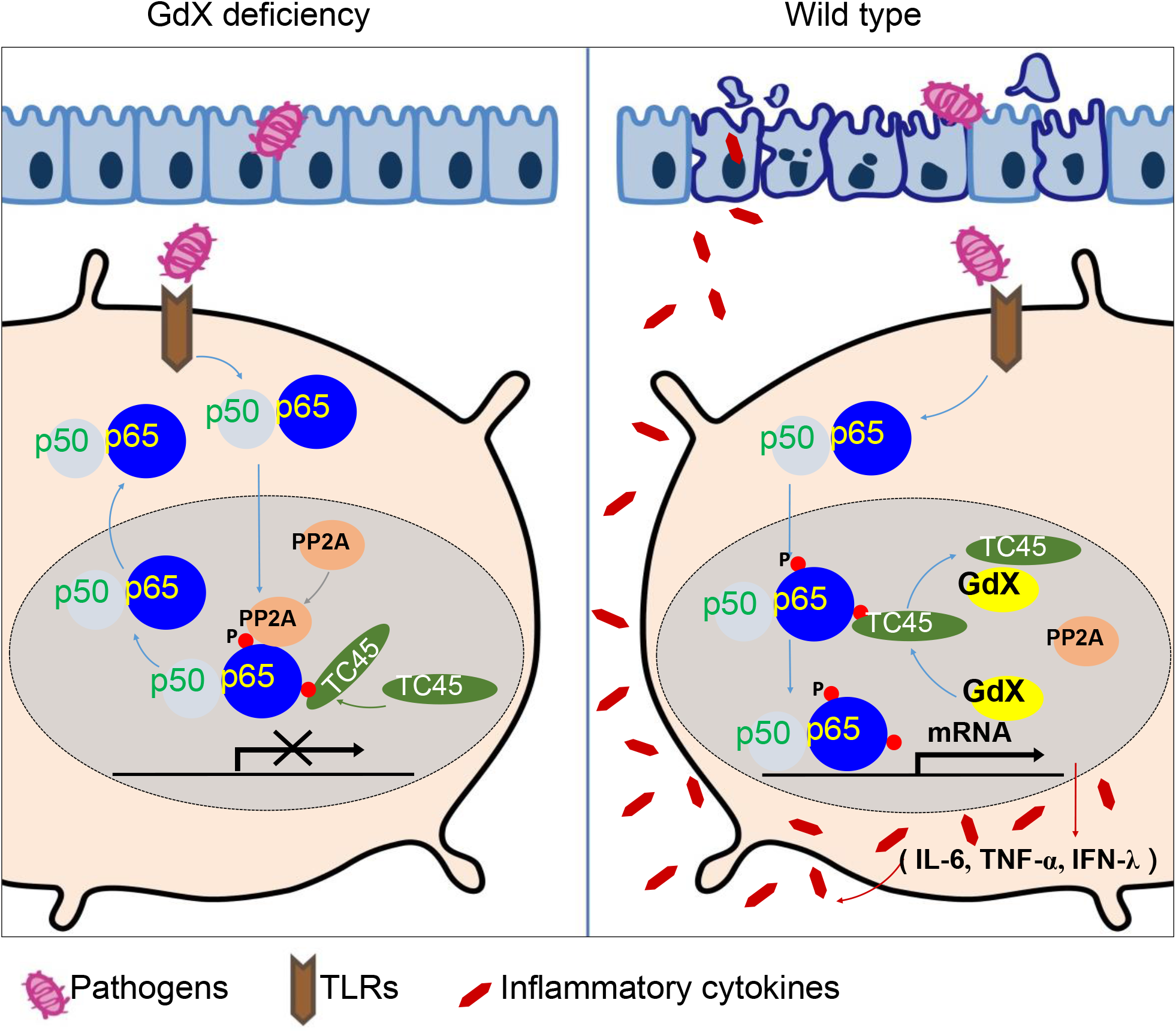
A model of the regulation of dephosphorylation of p65 by GdX. GdX maintained the level of p-p65 by blocking the complex formation of TC45 and PP2A with p65, leading to abrogated dephosphorylation of p65. Without GdX, TC45 brings PP2A to associate with p-p65 and mediates dephosphorylation of p-p65 for the termination of NF-kB signaling. In the presence of GdX, GdX interacts with TC45 and competes TC45 from interaction of PP2A and p-p65, leading to impaired dephosphorylation of p-p65. In such a way, GdX guards TC45 and PP2A from interaction with p65 to maintain DCs and Mφs alert to pathogen attacks.

Although NF-κB signaling is tightly associated with IBDs, the cell-type specific mechanisms of NF-κB is poorly understood. GdX deficiency led to reduced phosphorylation of p65 in colon sections during mucosal inflammation, which resulted in a less severe colitis. Consistent with our studies, the level of p65 has been reported increased in the nuclear extracts of intestinal lamina propria biopsy from IBD patients (Ardite et al., 1998). Moreover, activated p65 was found in either Mφ or epithelial cells from inflamed mucosa but was almost absent in normal mucosa (Roesky et al., 2000). We previously observed that IECs from GdX^−/Y^ mice had a greater proliferative ability which endowed an enhanced ability to maintain mucosal integrity (Wang et al., 2014). In this study, we found that deletion of GdX in IECs had no effect on DSS-induced colitis, but GdX^ΔDC^ and GdX^ΔMφ^ mice displayed alleviated mucosal inflammation. Hence, we conclude that GdX mainly functions in DCs and Mφ to regulate the acute colitis. This finding also implied that the NF-κB signaling in DCs and Mφ is critical for tissue inflammation and damage. This notion is supported by other studies that SCID mice developed intestinal inflammation similar to that developed in immunocompetent Balb/c mice after DSS-treatment (Tlaskalova-Hogenova et al., 2005), and adoptive transfer of GMDCs exacerbated DSS-induced colitis while ablation of DCs ameliorated the colitis (Berndt et al., 2007). Our results are also supported by the depletion of Mφ (Watanabe et al., 2003), inhibition of myeloid cells by an antibody against CD11b/CD18 (Mac-1) (Palmen et al., 1995) or deletion of IL-6 reduced intestinal inflammation (Naito et al., 2004). The function of GdX in DCs and Mφ in the regulation of DSS-induced colitis echoes the observations that DCs and Mφ (Coombes and Powrie, 2008) are direct contributors in response to acute mucosal damage during intestinal inflammation after DSS treatment (Tlaskalova-Hogenova et al., 2005).

p65 phosphorylation is critical for activation of NF-κB-dependent transcription (Sakurai et al., 1999; Sizemore et al., 2002; Zhong et al., 1998). Our data showed that GdX prolonged LPS-induced phosphorylation of p65 by trapping TC45 and PP2A. Consequently, p65 phosphorylation is sustained by GdX. Mechanistically, we observed that TC45 was critical for GdX-maintained p65 phosphorylation on serine 536. As dephosphorylation of serine 536 of p65 is induced by PP2A (Hsieh et al., 2011), we were considering that TC45 might function as a co-factor to bridge PP2A to p-p65, similar to the function of PHF20 (Zhang et al., 2013). However, TC45 failed to interact with PP2A. Therefore, we proposed a model the TC45 interacts with the N-terminus of p65 via Y100 and releases the coating of the C-terminus of p65 for PP2A association. Our functional results further demonstrated that mutant p65 (Y100F) impaired the interaction of TC45 to inhibit the transcriptional activity of p65. These results suggested that Y100 is critical for TC45 interaction and GdX functions to complete via Y100. However, further works are required to analyze the whole structure of p65, and how the interaction of TC45 with the N-terminus of p65 mediates the interaction of PP2A at the C-terminus of p65 remains to be elucidated.

In summary, our study provided the evidence that GdX promote the pro-inflammatory phenotypes of DCs and Mφ by activating NF-κB signaling. Targeting GdX in innate immune cells might offer therapeutic benefits for colitis, particularly for the development of DC- or Mφ-based therapeutic strategies for acute inflammation.

## Materials and methods

### Mice

C57BL/6J mice were housed in a specific pathogen-free (SPF) facility at Tsinghua University. GdX^−/Y^ and GdX^fl/fl^ mice were generated as described previously (Wang et al., 2012), and have been backcrossed to the C57BL/6J background for at least eight generations. Floxed GdX mice were bred with CD11c-Cre (Jackson stock 008068, B6.Cg-Tg (Itgax-cre) 1-1Reiz/J), LysM-Cre (Jackson stock 004781, B6.129P2-Lyz2^tm1(cre)Ifo^/J) or Villin-Cre (Jackson stock 018963, B6N.Cg-Tg (Vil-cre) 997Gum/J). Cre-negative littermates were used as controls.

### Assessment of the biological effects of LPS

To compare the survival rate under endotoxin shock, mice were injected with 30 mg/kg of LPS from Escherichia coli 055:B5 (Sigma, St. Louis, MO, USA). Mice were injected with 20 mg/kg LPS for induction of acute inflammation and serums were collected for cytokine measurement by using ELISA assay.

### RNA sequencing

The WT and GdX-deficient mice were injected (i.p.) with 20 mg/kg LPS, and then the splenocytes were collected after 1.5 or 6 h. RNAs were purified and reverted into cDNA libraries. High-throughput sequencing was performed by BGISEQ-500 (Beijing Genomics Institute, BGI). The RNA-seq was carried out with two biological replicates. The RNA-seq reads were mapped to mm10 genome by HISAT2 v2.0.4. The differentially expressed gene (DEG) reads were analyzed by DEGseq. The genes with absolute log2 >1 fold changes and a threshold q value<0.05 were regarded as DEGs. GO analysis was performed using differentially expressed genes against gene sets from DAVID GO database (https://david.ncifcrf.gov/) and KEGG pathway database (http://www.genome.jp/kegg/pathway.html). Sequencing data have been deposited in GEO under accession code GSE116956.

### Cell culture

For Fms-like tyrosine kinase 3 ligand (Flt3L)-supplemented BM-cultured DCs (FLDCs) preparation, bone marrow (BM) cells were cultured in RPMI-1640 complete medium (RPMI-1640-10% FBS-1% P/S) in the presence of recombinant murine Flt3L (200 ng/mL) for 7-8 days, and then sorted for pDCs, CD24^+^ cDCs and CD24^−^ cDCs. For granulocyte-macrophage colony-stimulating factor-induced DCs (GMDCs) preparation, BM cells were incubated with GM-CSF (20 ng/mL) and IL-4 (20 ng/mL) for 6-7 days. For bone marrow derived macrophages (BMDMs), BM cells were cultured in macrophage colony-stimulating factor (M-CSF) conditional medium (DMEM-10% FBS-1% P/S with the supplement of L929 cell supernatant) for 5-6 days. All of the cytokines were purchased from PeproTech (Rocky Hill, NJ, USA). For isolation of peritoneal Mφ, mice were injected with 4% thioglycollate medium in a total volume of 0.8 ml, and then peritoneal Mφ were isolated 3 days later.

### Cytokine secretion analyses

FLDCs were seeded in 96-well plates and treated with TLR agonists, including 50 nM Pam2CSK4 (InvivoGen, San Diego, CA, USA, tlrl-pm2s-1), 50 nM Pam3CSK4 (InvivoGen, tlrl-pms), 100 μg/ml Poly (I:C) (InvivoGen, tlrl-pic), 100 ng/mL LPS (Sigma), 1 μg/ml R848 (InvivoGen, tlrl-r848), 10nM ODN 1668 (AdipoGen, San Diego, CA, USA, IAX-200-001) and 1uM ODN 2216 (AdipoGen, IAX-200-005), respectively for 16 hrs. Supernatants were then collected for ELISA analyses of IL-6, TNF-α, IL-12p40, IL-12p70, MIP-1α, RANTES and IFN-λ respectively. Recombinant mouse RANTES (R&D, Minneapolis, MN, USA, 478-MR), MIP-1α (R&D, 450-MA), IFN-λ3 (R&D, 1789-ML), IL-12p70 (eBioscience, Palo Alto, CA, USA, 14-8121), TNF-α (eBio, 39-8321-65) and IL-6 (eBio, 39-8061-65) were used as internal standard protein controls. Antibodies against mouse RANTES (R&D, MAB4781), MIP-1α (R&D, AF-450-NA), IFN-λ2/3 (R&D, MAB17892), IL-12p70 (BioLegend, San Diego, CA, USA, 511802), IL-12/IL-23p40 (eBio, 14-7125-85), IL-6 (eBio, 14-7061-81) and IFN-α (PBL, 22100-1) were used as capture antibodies. Biotinylated monoclonal antibodies against mouse RANTES (R&D BAF478), MIP-1α (R&D BAF450), IFN-λ3 (R&D BAM17891), IL-12/IL-23p40 (eBio 13-7123), TNF-α (eBio14-7423-81), IL-6 (eBio13-7062-81) were used in combination with streptavidin-HRP (Amersham, Little Chalfont, UK, RPN4401) for ELLSA analyses.

### Transfection and luciferase reporter assay

HEK293T cells were transfected with plasmids encoding κB-luciferase reporter, pRL-TK Renilla luciferase, and different expression vectors using Lipofectamine-2000 (Invitrogen, Carlsbad, CA, USA). The κB-luciferase reporter activity was determined by a dual luciferase assay kit (Promega, Madison, WI, USA) as previously reported (Liu et al., 2015).

### Antibodies, immunoprecipitation and immunoblot analyses

Total cell lysates were prepared after transfection or stimulation for the experiments. For IP experiments, cell extracts were incubated with indicated antibodies together with Protein A/G beads (Pierce, Rockford, IL, USA) overnight. Beads were washed four times with lysis buffer, and immunoprecipitates were eluted with SDS loading buffer (Cell Signaling Technology, Danvers, MA, USA) and resolved in SDS-PAGE gels. Antibodies used were rabbit polyclonal antibodies against p65, p-p65, IkBα, p-IkBα, ERK, p-ERK, p38, p-p38, JNK, and p-JNK (from Cell Signaling) and mouse monoclonal antibodies against Myc (9E10), HA (Santa Cruz, Dallas, Texas, USA), Flag (M2, Sigma). Anti-GdX antibodies were generated in this lab, and the specificity was examined (Wang et al., 2012). WB and IP experiments were performed according to previous protocols (Liu et al., 2015).

### Quantitative RT-PCR

Total RNA was isolated with the RNeasy kit (Qiagen, Germantown, MD, USA), and cDNA was synthesized with SuperScript RT III (Invitrogen). The mRNA levels of IL-6, IL-1β, TNF-α, GdX and GAPDH were measured by real-time PCR performed in SYBR Green I on 7900 real-time PCR detection system (Applied Biosystems, Grand Island, NY, USA). Primer sequences were listed as follows: mouse IL-1β (forward: 5′-CTCCATGAGCTTTGTACAAGG -3′, reverse: 5′-TGCTGATGTACCAGTTGGGG -3′), mouse TNF-α (forward: 5′-CGGACTCCGCAAA GTCTAAG-3′, reverse: 5 ‘-ACGGCATGGATCTCAAAGAC-3′), mouse IL-6 (forward: 5′-GGAAATTGGGGTAGGAAGGA -3′, reverse: 5′-CCGGAGAGGAGACTTCACAG -3′), mouse GdX (forward: 5′-AGCACCTGGTCTCGGATAAG -3′, reverse: 5′-GCCCAATGTTGTAATCTGACAG-3′) mouse GAPDH (forward: 5′-TGTGTCCGTC GTGGATCTGA-3′, reverse: 5 ‘-CCTGCTTCACCACCTTCTTGA-3′). PCR was carried out for 35 cycles using the following conditions: denaturation at 95°C for 20 s, annealing at 58°C for 20 s, and elongation at 72°C for 20 s.

### Structural Analysis

All protein interacting models were predicted by Z-Dock (v.3.0.2)(Pierce et al., 2011). The proteins accession numbers used for docking models are 2ie3 (PP2A, 1-309), 2n22 (p65, 521-551), 1nfi (p65, 20-320), 1l8k (TC45, 1-314) and 2dzi (Gdx, 1-74), which were obtained from RCSB Protein Data Bank(Iversen et al., 2002; Jacobs and Harrison, 1998; Lecoq et al., 2017; Xing et al., 2006). The results of the models were visualized and processed by the PyMOL Molecular Graphics System (Version 1.8 Schrödinger, LLC).

### DSS-induced colitis

Mice were subjected to acceptance of 3% (wt/vol) DSS (molecular weight, 36,000–50,000; MP Biomedicals Santa Ana, CA, USA) in drinking water ad libitum for 6 days. Body weight and stool were monitored daily starting from day 0 of treatment. Colons were collected after mice were sacrificed, fixed in 10% (vol/vol) formaldehyde, and sectioned for immunohistological staining with indicated antibodies and for H&E staining according to protocols used previously (Wang et al., 2014).

### Statistical analyses

All of the *in vitro* experiments were repeated at least three times whereas all *in vivo* experiments were performed at least twice. All statistical analyses were calculated using GraphPad Prism 5. Data were presented as mean ± SEM. Statistical significance was determined with the two-tailed unpaired Student’s t test. Differences were considered to be statistically significant when p < 0.05.

## Acknowledgements

This work was supported by grants from the Chinese National Major Scientific Research Program (2015CB943200, 2013ZX08011-006, 2011CB910502), grants from the National Natural Science Foundation of China (81372167, 81301700, 81572728, 31330027, 81402293, 81372372), and the Tsinghua Science Foundation (20121080018, 20111080963). We thank Dr. Andrew Larner to provide TC45 KO cells. We thank Dr. Xinquan Wang and Dongli Wang for providing the assistance in the molecular docking analyses. We thank Taylor Chrisikos and Rachel Babcock for revising the manuscript.

## Supplement Information

### Figure legends

**Figure 1-figure supplement 1.**
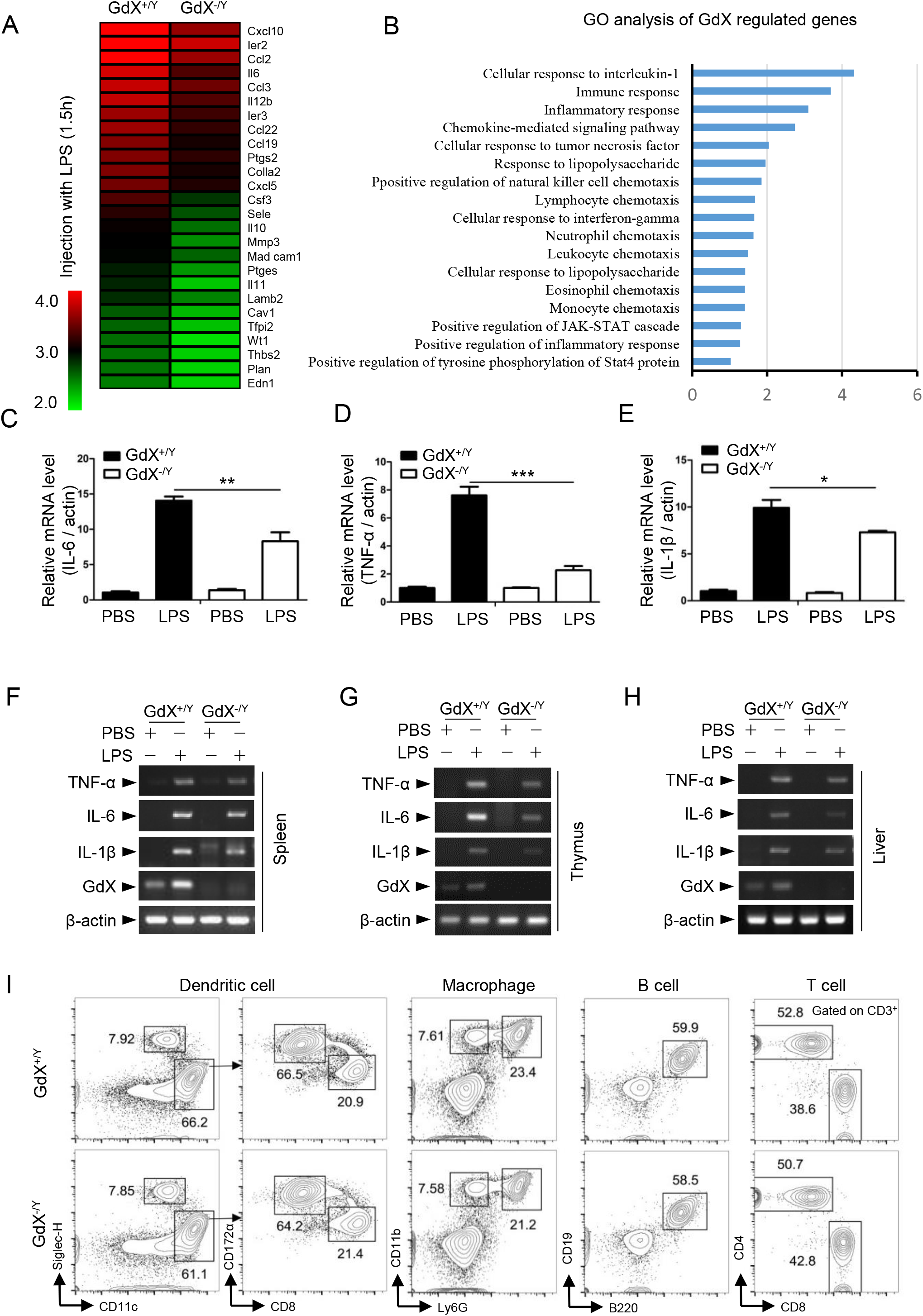
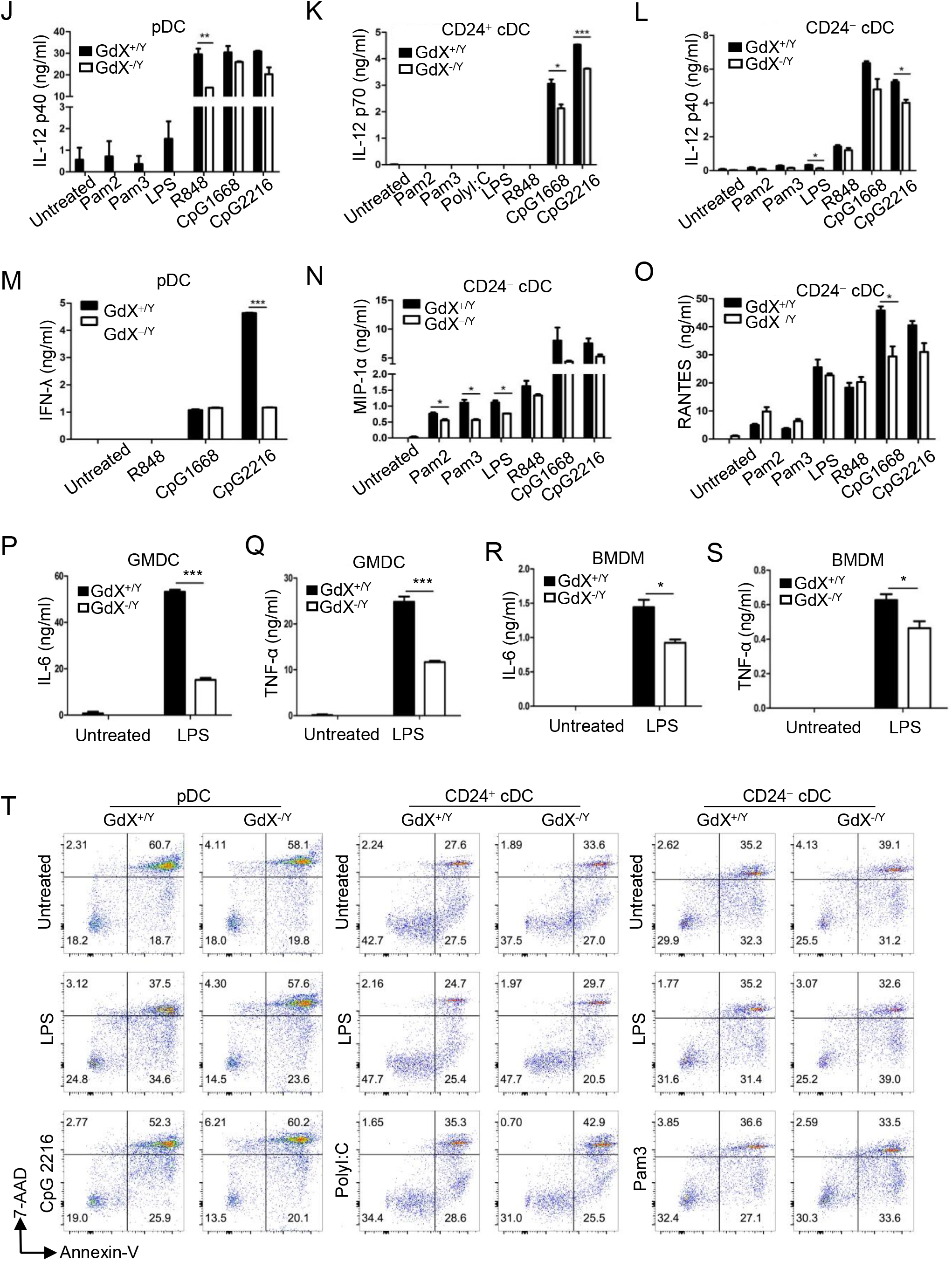
Decreased inflammatory cytokines were observed in the tissues of GdX^−/Y^ mice. (A) The heat map of the downregulated NF-κB target genes (list of 26 genes, absolute log2 >1, false discovery rate <0.05, p<0.05) in splenocytes of GdX^−/Y^ mice and wildtype littermates (n=4) injected with LPS. The mice were injected with LPS (20 mg/kg body weight, ip), and 1.5 hrs after injection, RNA was extracted from their splenocytes and performed the RNA-seq analysis. (B) GO term analysis of GdX function in the splenocytes of LPS-administrated mice showed one of the most significantly enriched biological process is related to inflammatory response. (C-E) Deletion of GdX down regulated the gene expression of pro-inflammatory cytokines in the spleen. Quantitative RT-PCR analyses were performed for the mRNA levels of IL-6 (C), TNF-α (D) and IL-1β (E) in the spleen from mice (n=3) treated with LPS. (F-H) mRNA levels of inflammatory cytokines and GdX in splenocytes (F), thymocytes (G) and hepatocytes (H) were analyzed by RT-PCR from GdX^+/Y^ and GdX^−/Y^ mice challenged with LPS (20 mg/kg) for 16 h. (I) The development of different immune cells was normal in GdX^−/Y^ mice. The lymphoid tissues were harvested from GdX^+/Y^ (n=6) and GdX^−/Y^ mice (n=6), and then analyzed the cell numbers of different immune cells and their progenitors. The progenitors were Lin^−^c-kit^hi^Sca-1+HSC (hematopoietic stem cell), Lin^−^c-kit^lo^Sca-1+CD127^+^ CLP (common lymphoid progenitor), Lin^−^ c-kit^hi^Sca-1^−^CD16/32^lo^CD34^+^ CMP (common myeloid progenitor), Linc-kit^hi^Sca-1^−^ CD16/32^hi^CD34^+^ GMP (granulocyte-macrophage progenitor) and Lin^−^c-kit^lo^CD135^+^CD115^+^CD11c^−^ CDP (common dendritic cell progenitor). (J-L) Different subtypes of FLDCs from GdX^+/Y^ and GdX^−/Y^ mice were used to measure the production of IL-12 in response to indicated TLR ligand stimulations. (M) IFN-λ production was decreased in GdX-deficient pDCs after stimulation by TLR agonists for 16 h. (N and O) Chemokines from GdX-deficient CD24^−^ cDCs were analyzed by ELISA after stimulation by TLR agonists for 16 h. The results were presented as mean ±SEM from three repeats. *, p < 0.05; **, p < 0.01; ***, p < 0.001. (P-S) Cytokine production (IL-6 and TNF-α) in supernatants of GMDCs and BMDMs after LPS stimulation (100 ng/mL) for 16 h were measured by ELISA. The results were presented as mean ± SEM from three repeats. *, p < 0.05; **, p < 0.01; ***, p < 0.001. (T) GdX deletion did not influence the apoptosis in DCs after stimulation by TLR agonists. pDCs, CD24^+^ cDCs, CD24^−^ cDCs were incubated with different TLR agonists for 16 h, and apoptosis was detected by FACS with 7-Amino-actinomycin D (7-AAD) and Annexin-V staining. Annexin-V^+^/7-AAD^−^ (early apoptosis) and Annexin-V^+^/7-AAD^+^ (late apoptosis) cells were quantified.

**Figure 2-figure supplement 1.**
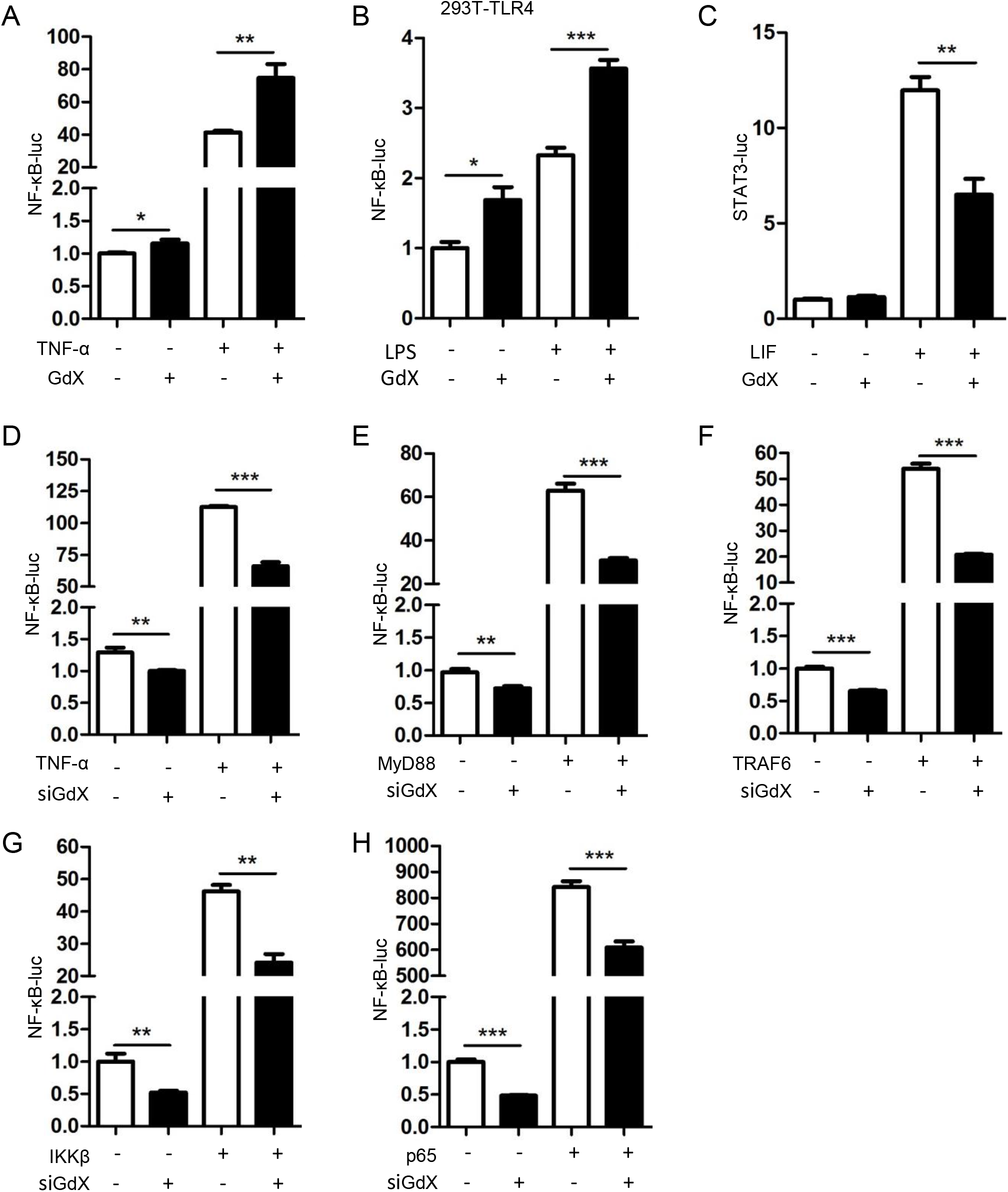
(A and B) GdX promoted the transcriptional activity of NF-κB. HEK293T cells, with or without over-expression of GdX, were co-transfected with NF-κB luciferase reporter (NF-κB-luc) and Renilla luciferase reporter. Relative luciferase activities were determined in three independent experiments after the cells were treated with TNF-α (10 ng/mL) (A) and LPS (100ng/ml) (B) for 8 h. (C) GdX inhibited STAT3 transcriptional activity. Luciferase activities were determined in HEK293T cells, transfected with the APRE-luc reporter treated with or without LIF. (D) Depletion of GdX decreased the activity of NF-κB. Relative luciferase activities were determined with or without depletion of GdX (siGdX, sequences targeting human GdX) after TNF-α (10 ng/mL) treatment. (E-H) Depletion of GdX decreased the activity of NF-κB. HEK293T cells were transfected with NF-κB-luc, together with MyD88 (E), TRAF6 (F), IKKβ (G), or p65 (H), along with or without GdXi. Luciferase activity was measured at 36 h after transfection and the results were presented as mean ± SD from three repeats. **, p < 0.01; ***, p < 0.001. (I) The level of p-p65 was significantly lower in splenocytes from GdX^−/Y^ mice than that from GdX^+/Y^ mice. Splenocytes were isolated and treated with TNF-α (10 ng/mL) for 15 min.

**Figure 3-figure supplement 1.**
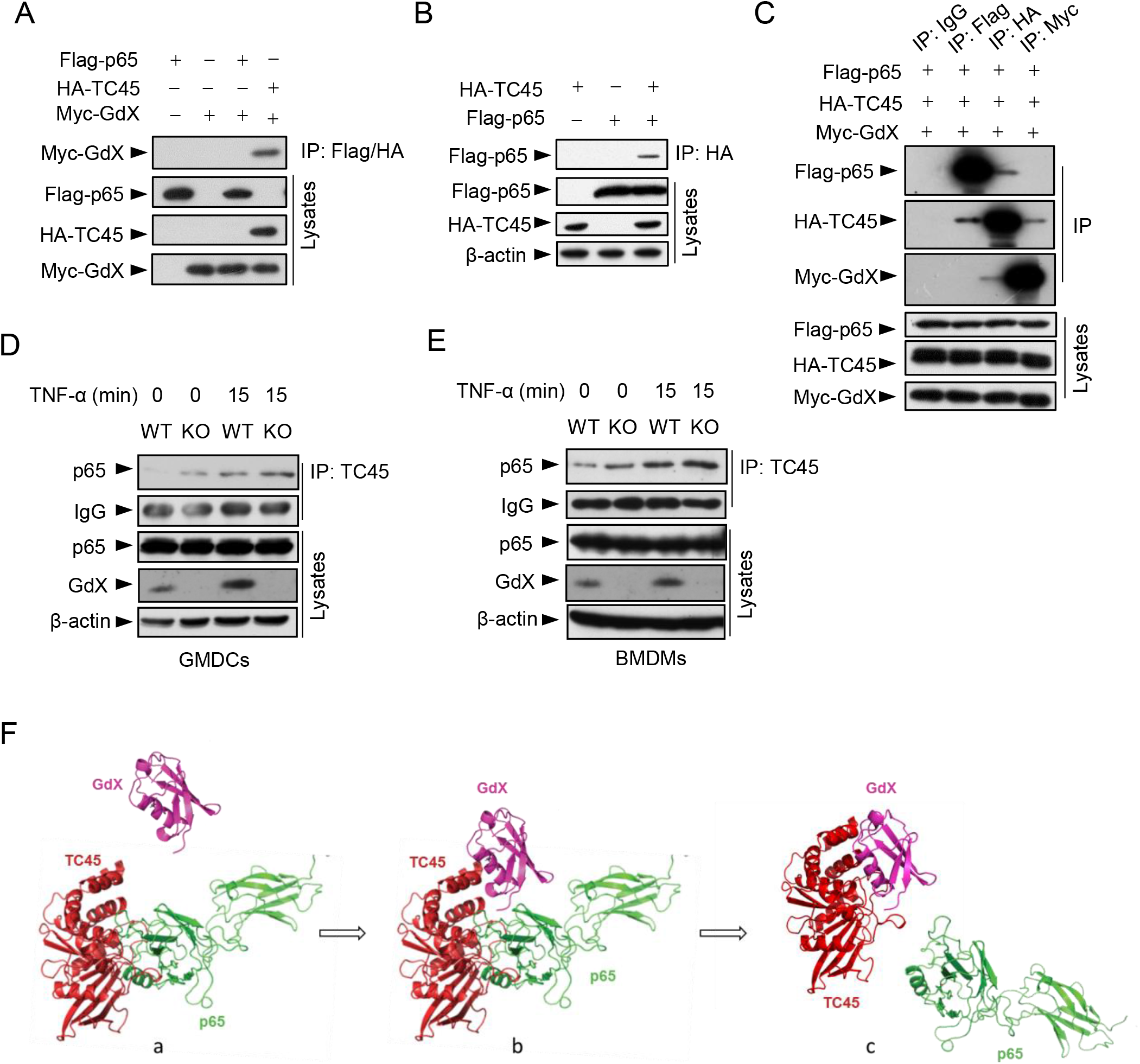
(A) GdX failed to interact with p65. HEK293T cells were transfected with the indicated plasmids and IP was performed with a mixture of an antibody against Flag and an antibody against HA. (B) HA-TC45 interacted with Flag-p65. An IP experiment was performed by using an anti-HA antibody. (C) TC45 interacted with either p65 or GdX. IP experiments were performed by using anti-Flag, anti-HA or anti-Myc antibody after HEK-293T cells were transfected with Flag-p65, HA-TC45 and Myc-GdX for 24 h. IgG was used as a negative control. This experiment indicated that two separate complexes Flag-p65/HA-TC45 and Myc-GdX/HA-TC45 formed. (D and E) The interaction of TC45 and p65 was increased in GdX-depleted (KO) GMDCs and BMDMs. GMDCs (D) and BMDMs (E) were used to perform the endogenous IP assay with TNF-α (10 ng/mL) treatment for 15 min. GMDCs and BMDMs from wildtype (WT) mice were used as controls. (F) A docking model of the competition of GdX with TC45 to interact with p65. While TC45 interacts with p65 (a), there are two helixes in TC45 which are free from interacting with p65 but are able to associate with GdX (b). In this way, GdX occupies the interface of TC45 to complete off p65 (c).

**Figure 4-figure supplement 1.**
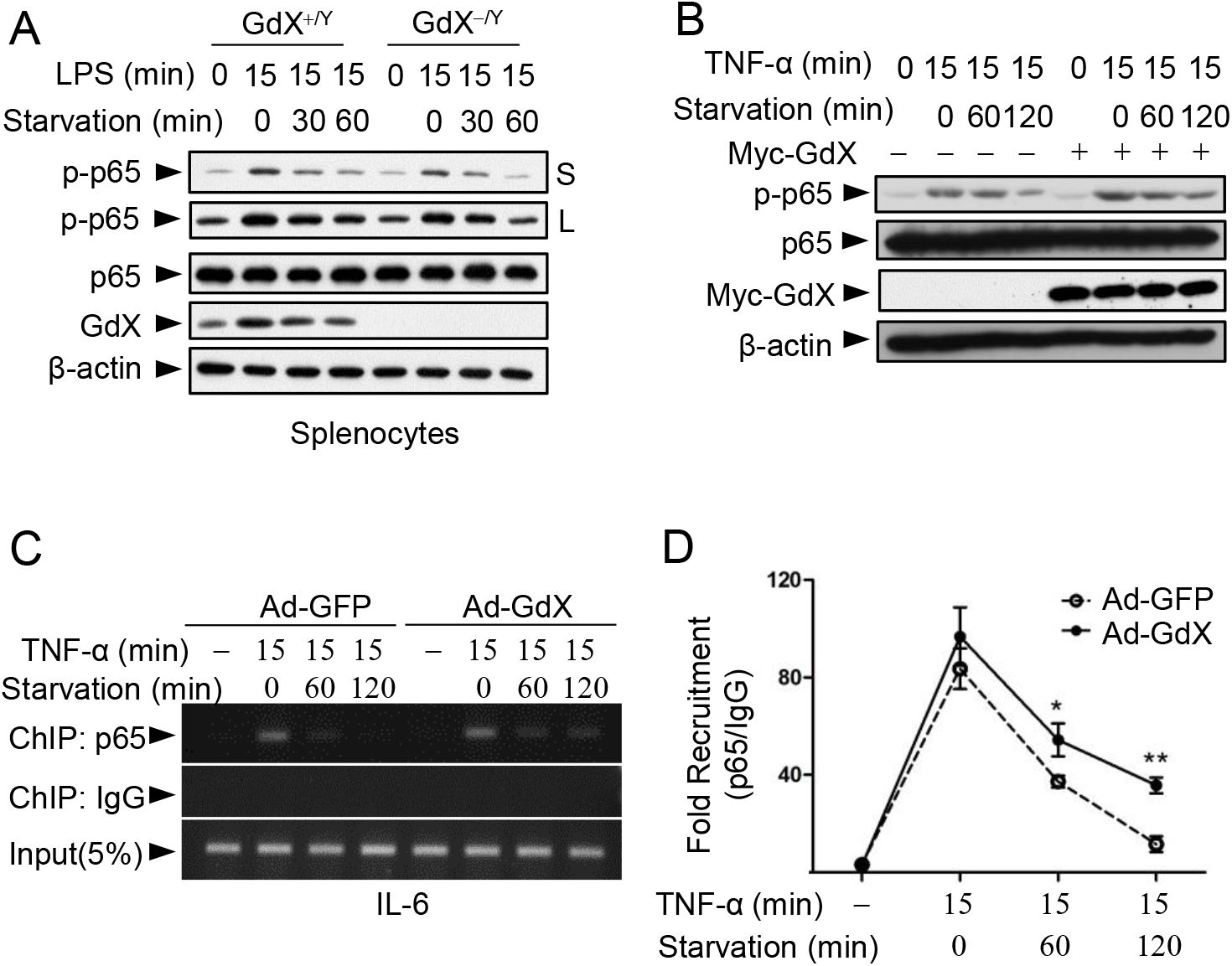
(A) The level of p-p65 was decreased in GdX-depleted splenocytes. Splenocytes were treated with LPS (100ng/ml) and then subjected to starvation for indicated times. Levels of p-p65 imaged at different exposure times (S, short; L, long) were showed. (B) Over-expression of GdX inhibited p65 dephosphorylation in response to TNF-α treatment. (C) GdX increased the DNA binding ability of p65. DC2.4 cells were infected by an adenovirus expressing GFP or GdX and were stimulated with TNF-α (10 ng/mL) for the 15 min, and then subjected to starvation. Chromatin was immunoprecipitated with an antibody against p65. PCR was performed by using primers targeting NF-κB binding sites on the *IL-6* gene promoter. Five percent of the precipitated chromatin was assayed to verify an equal loading (Input). (D) Real time PCR was used to quantify the amounts of chromatin-immunoprecipitated DNA from B.

**Figure 5-figure supplement 1.**
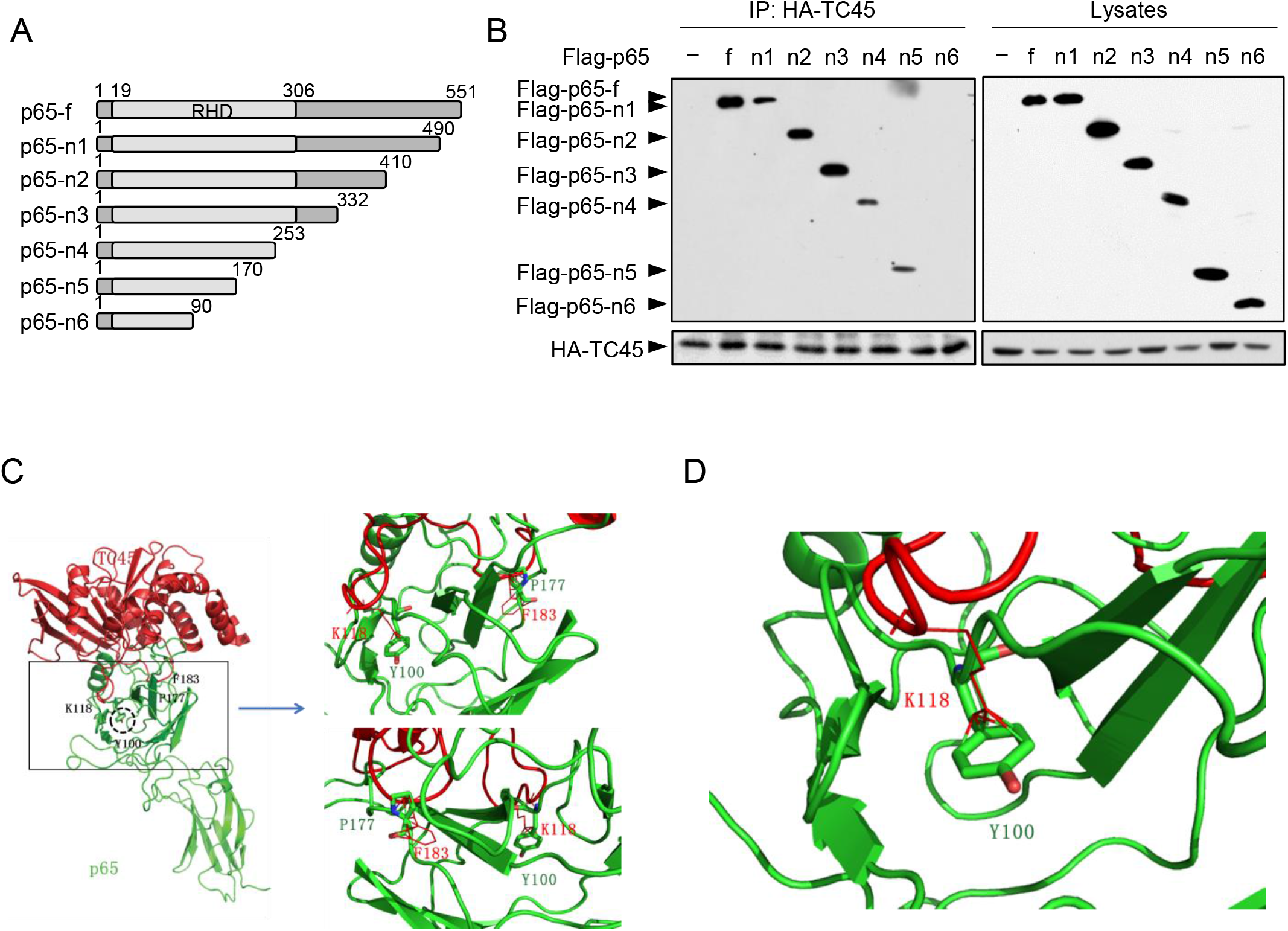
(A) A schematic diagram showing domain structures of p65 and its deletions. RHD: rel homology domain. Letter “f” indicated full length protein and different deletions were showed as n1 to n6. Numbers indicated the amino acid positions. (B) TC45 failed to interact with p65-n6. HEK293T cells were co-transfected with HA-TC45 and Flag-p65 or its deletions as indicated. Flag-tagged proteins were immunoprecipitated with an antibody against HA, and then the complex was blotted with an antibody against Flag. (C and D) A molecular docking analysis showed Y100 is critical for the interaction of p65 with TC45. (C) The interface of p65 and TC45 is maintained by F183 and K118 in TC45 and Y100 and P177 in p65. Two reviews are showed on the right panel. (D) A critical band between Y100 at p65 and K118 at TC45 is showed.

**Figure 6-figure supplement 1.**
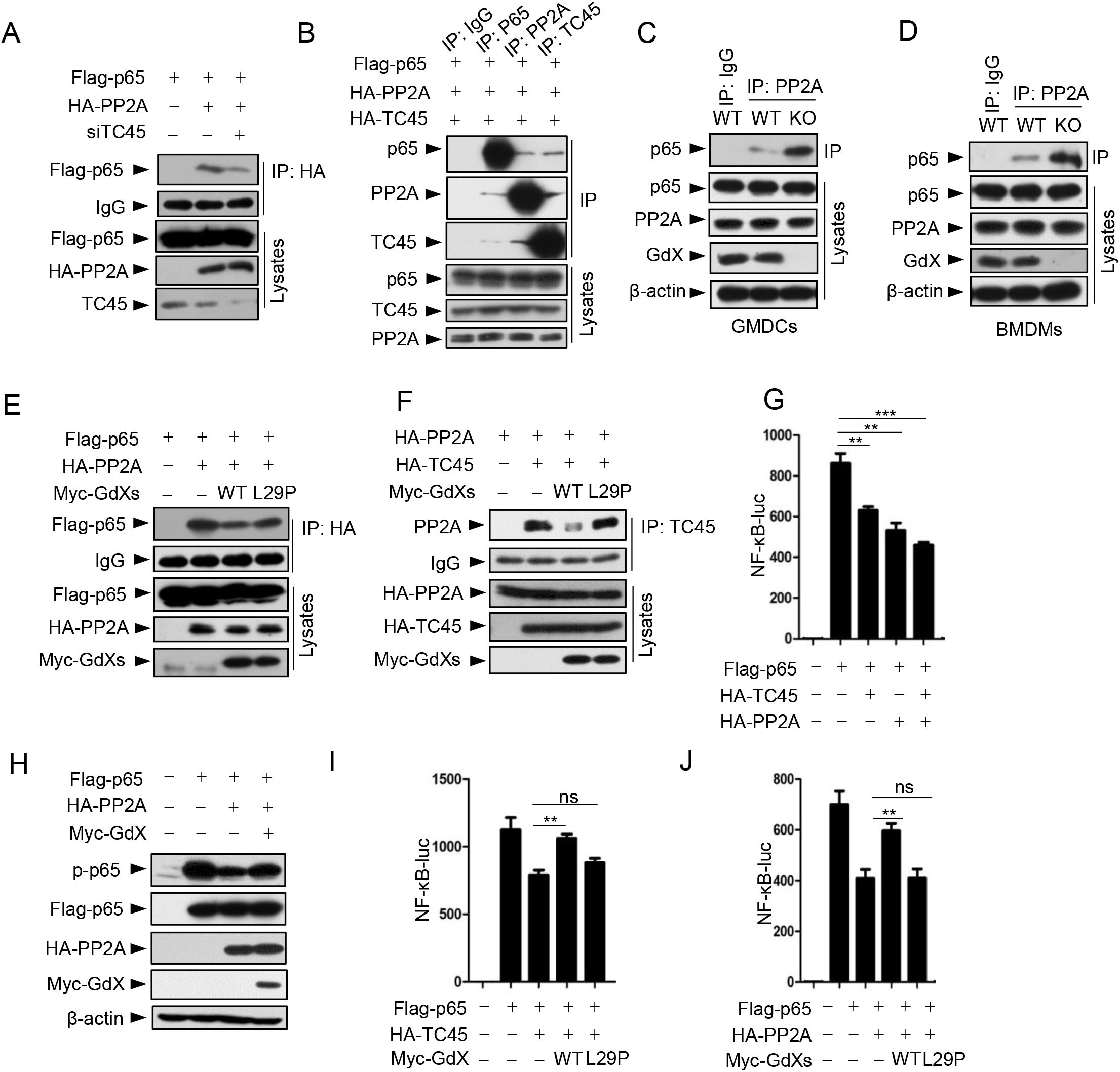
(A) The interaction of p65 and PP2A was decreased when TC45 was knock down. TC45 was depleted by transfection with a siRNA targeting TC45 (siTC45) in HEK293T cells for 36 h before harvesting the cells. An IP was performed using an antibody against HA. (B) p65, PP2A and TC45 formed a complex. IP experiments were performed by using anti-p65, anti-PP2A or anti-TC45 antibody after HEK293T cells were transfected with Flag-p65, HA-PP2A and HA-TC45 for 24 h. (C and D) The interaction of PP2A and p65 was increased in GdX-depleted cells. GMDCs (C) and BMDMs (D) derived from GdX^+/Y^ and GdX^−/Y^ mice were used for IP experiments. (E) GdX(L29P) mutant failed to decrease the interaction of PP2A and p65. HEK293T cells were transfected with the indicated plasmids before the IP experiment. (F) GdX(L29P) mutant failed to decrease the interaction of PP2A and TC45. (G) PP2A and TC45 synergistically inhibited the transcriptional activity of NF-κB. (H) GdX rescued the PP2A-mediated dephosphorylation of p-p65. HEK293T cells were transfected with the indicated plasmids. The protein expression levels were examined by Western blot. (I) GdX rescued the TC45-induced the inhibition of transcriptional activity of NF-κB. (J) GdX rescued the PP2A-induced the inhibition of transcriptional activity of NF-κB. Luciferase activity was measured at 36 h after transfection with the indicated plasmids and the results were presented as mean ± SD from three repeats. **, p < 0.01; ***, p < 0.001.

**Figure 7-figure supplement 1.**
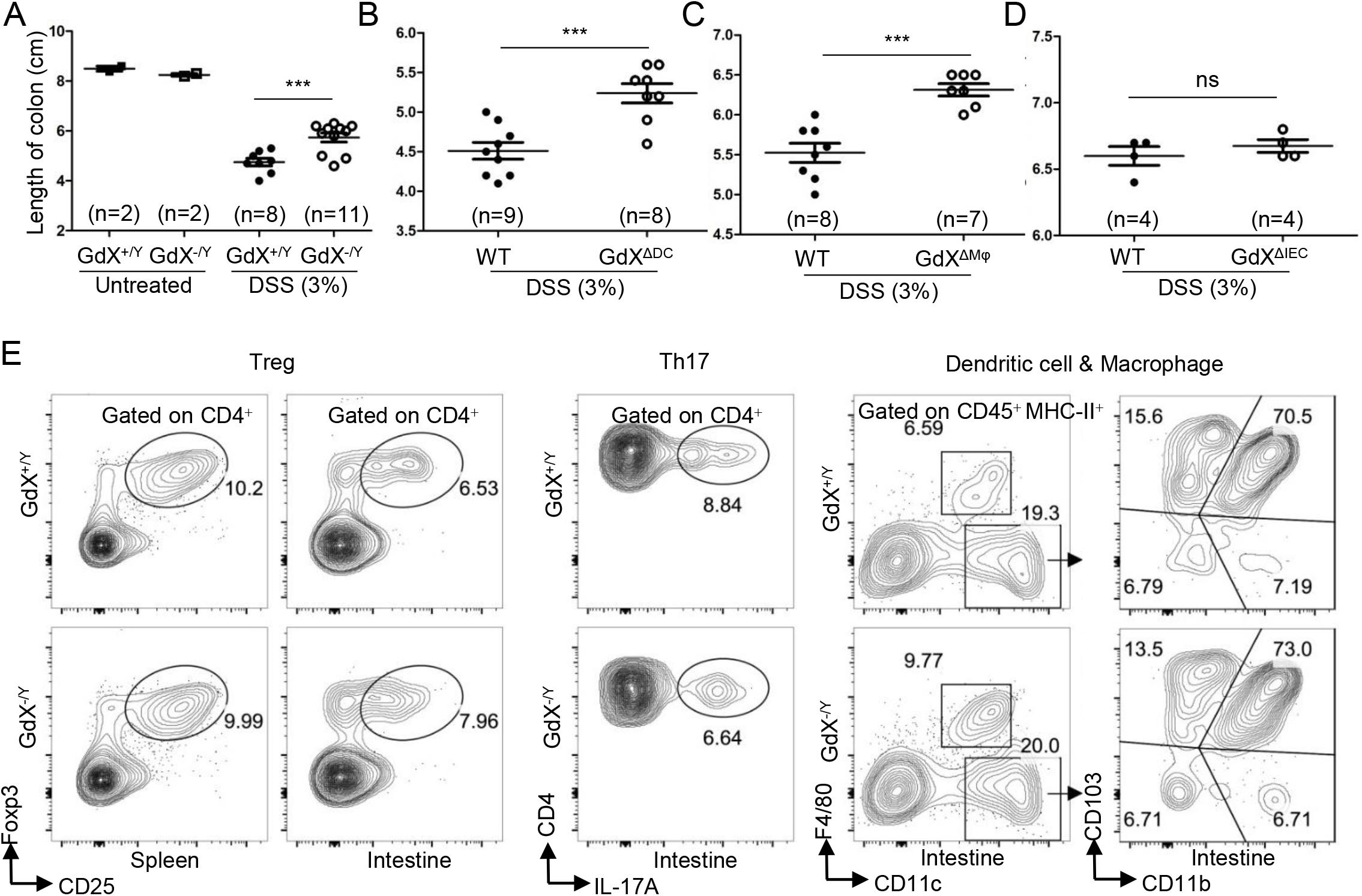
(A-D) The length of colon was determined after DSS treatment. The length of GdX^−/Y^ (A), GdX^ΔDC^ (B), GdX^ΔMφ^ (C) and GdX^ΔIEC^ (D) mice were compared with their WT (Cre-negative) littermates. Results were presented as means ± SD. The numbers of mice in each group is labeled. **P < 0.005. (E) Intestinal immune cell populations in GdX^ΔDC^ mice were similar to WT mice during DSS colitis. GdX^ΔDC^ mice and WT littermate were treated by 3% DSS for 6 day, and then the intestinal immune cells were purified. Splenic and intestinal T_reg_, intestinal Th17, intestinal F4/80^−^CD11c^hi^ DCs (including CD103^+^CD11b^−^, CD103^+^CD11b^+^ and CD103^−^CD11b^+^ DCs) and F4/80+CD11c^int^ Mφs were analyzed by FACS.

### Supplemental Experimental Procedures

#### Isolation of cells from tissues

For isolation DCs from spleen and thymus, tissue was minced with scissors and digested with 0.1 mg/ml DNaseI (Roche Molecular Biochemicals) and 1 mg/ml collagenase III (Worthington Biochemical) at 37°C for 25 min. Then, light-density cells were isolated in 1.077 g/cm^3^ (spleen) or 1.076 g/cm^3^ (thymus) Nycodenz (Axis-Shield) medium by centrifugation for 10 min at 1700 g. Additionally, splenocytes were incubated with mAb against CD3, CD90, TER119, Ly6G and CD19, followed by removal of non DC using anti-immunoglobulin (Ig)-coated magnetic beads (Bangs Laboratories). The enriched cells were stained with DC-specific markers and sorted.

For isolation lymphocytes from intestine lamina propria, the small intestine was taken out and removed off the mesentery, Peyer’s patches, fat and content. The small intestine was then moved into the medium (RPMI 1640, 1% P/S, 5 mM EDTA, 20 mM HEPES) and shook in 37°C incubator at 190 rpm for 30 min to wash off the epithelial cells. The remaining tissue was minced and digested with 10 U/ml collagenase CLISPA (Worthington Biochemical) and 0.1 mg/ml DNaseI at 37°C for 40 min. Subsequently, heavy-density cells were purified in 40% Percoll (GE Healthcare) by centrifugation for 10 min at 800 g.

For isolation of BM progenitors, the BM cells were depleted of red blood cells and followed by light-density separation (1.086 g/cm^3^ Nycodenz, 1700 g, 10 min) and then immune-magnetic bead negative selection (BM lineage cocktail: CD2, CD3, CD8, B220, CD11b, TER119, Ly6G).

Other immune cells were isolated from spleen, thymus and BM after removal of red blood cells.

#### Flow cytometry and antibodies

Single-cell suspensions were prepared and blocked in rat immunoglobulin (Jackson Laboratories) for 10 min for flow cytometry analyses. For cell surface markers, antibody incubation was performed at 4°C for 30 min. For intracellular Foxp3 staining, the Foxp3 Staining Buffer Set (eBioscience) was used. For intracellular IL-17A and TLR9 staining, we used Fixation /Permeabilization solution kit (BD). The following mAbs were used for cell staining and sorting: PE-Cy7-conjugated CD11c (N418), CD45R (RA3-6B2) and Ly6A/E (D7), PE-conjugated Siglec-H (eBio440c), F4/80 (BM8), CD3e (eBio500A2), CD135 (A2F10), Foxp3 (NRRF-30), IL-17A (eBio17B7), TLR2 (6C2) and CD103 (2E7), FITC-conjugated CD172α (P84), Ly6G (1A8), CD4 (GK1.5), CD16/32 (2.4G2), CD11c (N418), CD45R (RA3-6B2) and TLR9 (M9.D6), APC-conjugated CD8α (53-6.7), CD11b (M1/70), CD19 (eBio103), CD117 (ACK2), CD25 (PC-61.5.3), F4/80 (BM8) and CD24 (M1/69), eFluor 450-conjugated CD8α (53-6.7) and MHC-II (M5/114.15.2), APC-cy7-conjugated CD45 (30-F11), BV605-conjugated CD11b (M1/70), biotinylated CD127 (A7R34), CD34 (RAM34) and CD115 (AFS98). PE-streptavidin and PE-Cy7-streptavidin were used for second-stage staining. All of the antibodies were purchased from eBioscience, BD Biosciences or BioLegend. Dead cells were discriminated in all experiments using 7-AAD (eBio) staining. Cell apoptosis was analyzed by AnnexinV apoptosis detection kit (eBio). Cell analysis was carried out on LSRII and LSRFortessa flow cytometers; cell sorting was performed on FACSAria II and FACSAria III instruments. All of the machines were purchased form BD Bioscience. The purity of sorted populations was routinely more than 95%. Data analysis was performed on the single, live cell gate using FlowJo software (TreeStar).

#### Antigen presentation assay

The OT-I CD8^+^ and OT-II CD4^+^ T cells were isolated from the spleen of OT-I or OT-II transgenic mice, through depletion of red blood cells and immune-magnetic bead negative selection (CD8^+^ T cell cocktail: CD11b, F4/80, B220, CD11b, CD19, TER119, Ly6G, MHC-II,CD4; CD4^+^ T cell cocktail: CD11b, F4/80, B220, CD11b, CD19, TER119, Ly6G, MHC-II,CD8). And then the T cells were labeled using CFSE cell proliferation kit (Invitrogen).

Splenic CD8^+^ cDCs and CD8^−^ cDCs (1×10^5^ cells/ml) were sorted and incubated in 24-well plates (Costar-Corning) with OVA protein (100 ug/ml, Sigma) or OVA peptide (CD8^+^ cDCs: OVA257-264, 1 ng/ml; CD8^−^ cDCs: OVA peptide323-339, 10 ug/ml, Sigma) at 37 °C in RPMI 1640 complete medium. After 2 h, cells were washed twice. DC populations were plated with different numbers in 96-well round-bottom plates. Ten thousand CFSE-labeled OT-I or OT-II T cells (OT-I CD8^+^ T cells: CD8^+^ cDCs; OT-II CD4^+^ T cells: CD8^−^ cDCs) were added in each well in RPMI-1640 complete medium supplemented with 20 ng/ml GM-CSF. Proliferation was analyzed by flow cytometry after 60–90 h of culture.

#### Chromatin Immunoprecipitation (ChIP) Assay

A modified protocol from Upstate Biotechnology was used. Briefly, cells were fixed at 37°C for 10 min with 1% formaldehyde for crosslinking. The cells were resuspended in 500 ul of ChIP lysis buffer and mixed at 4°C and then sonicated for 30 s at level 2 (Ultrasonic Processor, Sonics) to yield DNA fragments that were 100–500 bp in size. Eluted DNA was recovered with QIAquick columns (Qiagen, Germany) and used as templates for PCR amplifications. The input control was from the supernatant before precipitation. The fragment corresponding to the NF-κB binding site in the IL-6 promoter was amplified by PCR with primers 5’-TGCTCAAGTGCTGAGTCACT-3’ and 5’-AGACTCATGGGAAAATCCCA-3’. Real time PCR was used to quantify the precipitated DNA fragments.

#### Immunofluorescent Analysis

Hela cells were plated on glass coverslips in 6-well dishes, incubated overnight at 37°C, and then infected with adenovirus. 24 h after infection, cells were rinsed with PBS three times, fixed with 4% paraformaldehyde in PBS for 15–20 min at room temperature, and permeabilized with 0.2% Triton X-100 in PBS for 10 min. Cells were blocked with 10% goat serum for 1 h at room temperature. The primary antibodies, diluted in PBS with 3% bovine serum albumin, were incubated overnight at 4°C, and bound antibodies were detected with secondary antibodies conjugated with TRITC (red) for 1 h at room temperature. Finally, cells were stained with DAPI. Stained cells were analyzed with a laser scanning confocal microscopy (OLYMPUS FV10i-Oil).

## References

Aderem, A., and R.J. Ulevitch. 2000. Toll-like receptors in the induction of the innate immune response. Nature 406:782–787.

Ahmad, S.F., M.A. Ansari, K.M. Zoheir, S.A. Bakheet, H.M. Korashy, A. Nadeem, A.E. Ashour, and S.M. Attia. 2015. Regulation of TNF-alpha and NF-kappaB activation through the JAK/STAT signaling pathway downstream of histamine 4 receptor in a rat model of LPS-induced joint inflammation. Immunobiology 220:889–898.

Ardite, E., J. Panes, M. Miranda, A. Salas, J.I. Elizalde, M. Sans, Y. Arce, J.M. Bordas, J.C. Fernandez-Checa, and J.M. Pique. 1998. Effects of steroid treatment on activation of nuclear factor kappaB in patients with inflammatory bowel disease. British journal of pharmacology 124:431–433.

Berndt, B.E., M. Zhang, G.H. Chen, G.B. Huffnagle, and J.Y. Kao. 2007. The role of dendritic cells in the development of acute dextran sulfate sodium colitis. J Immunol 179:6255–6262.

Chew, J., S. Biswas, S. Shreeram, M. Humaidi, E.T. Wong, M.K. Dhillion, H. Teo, A. Hazra, C.C. Fang, E. Lopez-Collazo, D.V. Bulavin, and V. Tergaonkar. 2009. WIP1 phosphatase is a negative regulator of NF-kappaB signalling. Nature cell biology 11:659–666.

Coombes, J.L., and F. Powrie. 2008. Dendritic cells in intestinal immune regulation. Nat Rev Immunol 8:435–446.

Ghosh, S., and M.S. Hayden. 2008. New regulators of NF-kappaB in inflammation. Nat Rev Immunol 8:837–848.

Hayden, M.S., and S. Ghosh. 2008. Shared principles in NF-kappaB signaling. Cell 132:344–362.

Hoesel, B., and J.A. Schmid. 2013. The complexity of NF-kappaB signaling in inflammation and cancer. Molecular cancer 12:86.

Hsieh, C.Y., G. Hsiao, M.J. Hsu, Y.H. Wang, and J.R. Sheu. 2014. PMC, a potent hydrophilic alpha-tocopherol derivative, inhibits NF-kappaB activation via PP2A but not IkappaBalpha-dependent signals in vascular smooth muscle cells. Journal of cellular and molecular medicine 18:1278–1289.

Hsieh, C.Y., M.J. Hsu, G. Hsiao, Y.H. Wang, C.W. Huang, S.W. Chen, T. Jayakumar, P.T. Chiu, Y.H. Chiu, and J.R. Sheu. 2011. Andrographolide enhances nuclear factor-kappaB subunit p65 Ser536 dephosphorylation through activation of protein phosphatase 2A in vascular smooth muscle cells. The Journal of biological chemistry 286:5942–5955.

Iversen, L.F., K.B. Moller, A.K. Pedersen, G.H. Peters, A.S. Petersen, H.S. Andersen, S. Branner, S.B. Mortensen, and N.P. Moller. 2002. Structure determination of T cell protein-tyrosine phosphatase. The Journal of biological chemistry 277:19982–19990.

Iwasaki, A. 2007. Mucosal dendritic cells. Annu Rev Immunol 25:381–418.

Jacobs, M.D., and S.C. Harrison. 1998. Structure of an IkappaBalpha/NF-kappaB complex. Cell 95:749–758.

Kawai, T., and S. Akira. 2011. Toll-like receptors and their crosstalk with other innate receptors in infection and immunity. Immunity 34:637–650.

Lecoq, L., L. Raiola, P.R. Chabot, N. Cyr, G. Arseneault, P. Legault, and J.G. Omichinski. 2017. Structural characterization of interactions between transactivation domain 1 of the p65 subunit of NF-kappaB and transcription regulatory factors. Nucleic acids research 45:5564–5576.

Li, H.Y., H. Liu, C.H. Wang, J.Y. Zhang, J.H. Man, Y.F. Gao, P.J. Zhang, W.H. Li, J. Zhao, X. Pan, T. Zhou, W.L. Gong, A.L. Li, and X.M. Zhang. 2008. Deactivation of the kinase IKK by CUEDC2 through recruitment of the phosphatase PP1. Nature immunology 9:533–541.

Li, S., L. Wang, M.A. Berman, Y. Zhang, and M.E. Dorf. 2006. RNAi screen in mouse astrocytes identifies phosphatases that regulate NF-kappaB signaling. Mol Cell 24:497–509.

Liu, C., Y. Zhang, J. Li, Y. Wang, F. Ren, Y. Zhou, Y. Wu, Y. Feng, Y. Zhou, F. Su, B. Jia, D. Wang, and Z. Chang. 2015. p15RS/RPRD1A (p15INK4b-related sequence/regulation of nuclear pre-mRNA domain-containing protein 1A) interacts with HDAC2 in inhibition of the Wnt/beta-catenin signaling pathway. The Journal of biological chemistry 290:9701–9713.

London, N.R., W. Zhu, F.A. Bozza, M.C. Smith, D.M. Greif, L.K. Sorensen, L. Chen, Y. Kaminoh, A.C. Chan, S.F. Passi, C.W. Day, D.L. Barnard, G.A. Zimmerman, M.A. Krasnow, and D.Y. Li. 2010. Targeting Robo4-dependent Slit signaling to survive the cytokine storm in sepsis and influenza. Science translational medicine 2:23ra19.

Molodecky, N.A., I.S. Soon, D.M. Rabi, W.A. Ghali, M. Ferris, G. Chernoff, E.I. Benchimol, R. Panaccione, S. Ghosh, H.W. Barkema, and G.G. Kaplan. 2012. Increasing incidence and prevalence of the inflammatory bowel diseases with time, based on systematic review. Gastroenterology 142:46–54 e42; quiz e30.

Murano, M., K. Maemura, I. Hirata, K. Toshina, T. Nishikawa, N. Hamamoto, S. Sasaki, O. Saitoh, and K. Katsu. 2000. Therapeutic effect of intracolonically administered nuclear factor kappa B (p65) antisense oligonucleotide on mouse dextran sulphate sodium (DSS)-induced colitis. Clinical and experimental immunology 120:51–58.

Naito, Y., T. Takagi, K. Uchiyama, M. Kuroda, S. Kokura, H. Ichikawa, R. Yanagisawa, K. Inoue, H. Takano, M. Satoh, N. Yoshida, T. Okanoue, and T. Yoshikawa. 2004. Reduced intestinal inflammation induced by dextran sodium sulfate in interleukin-6-deficient mice. International journal of molecular medicine 14:191–196.

Nathan, C. 2002. Points of control in inflammation. Nature 420:846–852.

Oeckinghaus, A., and S. Ghosh. 2009. The NF-kappaB family of transcription factors and its regulation. Cold Spring Harbor perspectives in biology 1:a000034.

Palmen, M.J., C.D. Dijkstra, M.B. van der Ende, A.S. Pena, and E.P. van Rees. 1995. Anti-CD11b/CD18 antibodies reduce inflammation in acute colitis in rats. Clinical and experimental immunology 101:351–356.

Pierce, B.G., Y. Hourai, and Z. Weng. 2011. Accelerating protein docking in ZDOCK using an advanced 3D convolution library. PloS one 6:e24657.

Roesky, H.W., A. Stasch, H. Hatop, C. Rennekamp, H.H. D, M. Noltemeyer, and H.G. Schmidt. 2000. A Facile Route to Group 13 Difluorodiorganometalates. Angewandte Chemie 39:171–173.

Sakurai, H., H. Chiba, H. Miyoshi, T. Sugita, and W. Toriumi. 1999. IkappaB kinases phosphorylate NF-kappaB p65 subunit on serine 536 in the transactivation domain. The Journal of biological chemistry 274:30353–30356.

Shu, G., L. Zhang, S. Jiang, Z. Cheng, G. Wang, X. Huang, and X. Yang. 2016. Isoliensinine induces dephosphorylation of NF-kB p65 subunit at Ser536 via a PP2A-dependent mechanism in hepatocellular carcinoma cells: roles of impairing PP2A/I2PP2A interaction. Oncotarget

Sizemore, N., N. Lerner, N. Dombrowski, H. Sakurai, and G.R. Stark. 2002. Distinct roles of the Ikappa B kinase alpha and beta subunits in liberating nuclear factor kappa B (NF-kappa B) from Ikappa B and in phosphorylating the p65 subunit of NF-kappa B. The Journal of biological chemistry 277:3863–3869.

Sugimoto, N., T. Rui, M. Yang, S. Bharwani, O. Handa, N. Yoshida, T. Yoshikawa, and P.R. Kvietys. 2008. Points of control exerted along the macrophage-endothelial cell-polymorphonuclear neutrophil axis by PECAM-1 in the innate immune response of acute colonic inflammation. J Immunol 181:2145–2154.

Tlaskalova-Hogenova, H., L. Tuckova, R. Stepankova, T. Hudcovic, L. Palova-Jelinkova, H. Kozakova, P. Rossmann, D. Sanchez, J. Cinova, T. Hrncir, M. Kverka, L. Frolova, H. Uhlig, F. Powrie, and P. Bland. 2005. Involvement of innate immunity in the development of inflammatory and autoimmune diseases. Annals of the New York Academy of Sciences 1051:787–798.

Toniolo, D., M. Persico, and M. Alcalay. 1988. A "housekeeping" gene on the X chromosome encodes a protein similar to ubiquitin. Proc Natl Acad Sci U S A 85:851–855.

Wang, Y., H. Ning, F. Ren, Y. Zhang, Y. Rong, F. Su, C. Cai, Z. Jin, Z. Li, X. Gong, Y. Zhai, D. Wang, B. Jia, Y. Qiu, Y. Tomita, J.J. Sung, J. Yu, D.M. Irwin, X. Yang, X. Fu, Y.E. Chin, and Z. Chang. 2014. GdX/UBL4A specifically stabilizes the TC45/STAT3 association and promotes dephosphorylation of STAT3 to repress tumorigenesis. Mol Cell 53:752–765.

Wang, Y., D. Wang, F. Ren, Y. Zhang, F. Lin, N. Hou, X. Cheng, P. Zhang, B. Jia, X. Yang, and Z. Chang. 2012. Generation of mice with conditional null allele for GdX/Ubl4A. Genesis 50:534–542.

Watanabe, N., K. Ikuta, K. Okazaki, H. Nakase, Y. Tabata, M. Matsuura, H. Tamaki, C. Kawanami, T. Honjo, and T. Chiba. 2003. Elimination of local macrophages in intestine prevents chronic colitis in interleukin-10-deficient mice. Digestive diseases and sciences 48:408–414.

Xing, Y., Y. Xu, Y. Chen, P.D. Jeffrey, Y. Chao, Z. Lin, Z. Li, S. Strack, J.B. Stock, and Y. Shi. 2006. Structure of protein phosphatase 2A core enzyme bound to tumor-inducing toxins. Cell 127:341–353.

Xu, Y., M. Cai, Y. Yang, L. Huang, and Y. Ye. 2012. SGTA recognizes a noncanonical ubiquitin-like domain in the Bag6-Ubl4A-Trc35 complex to promote endoplasmic reticulum-associated degradation. Cell reports 2:1633–1644.

Xu, Y., Y. Liu, J.G. Lee, and Y. Ye. 2013. A ubiquitin-like domain recruits an oligomeric chaperone to a retrotranslocation complex in endoplasmic reticulum-associated degradation. The Journal of biological chemistry 288:18068–18076.

Yang, J., G.H. Fan, B.E. Wadzinski, H. Sakurai, and A. Richmond. 2001. Protein phosphatase 2A interacts with and directly dephosphorylates RelA. The Journal of biological chemistry 276:47828–47833.

Zhang, T., K.A. Park, Y. Li, H.S. Byun, J. Jeon, Y. Lee, J.H. Hong, J.M. Kim, S.M. Huang, S.W. Choi, S.H. Kim, K.C. Sohn, H. Ro, J.H. Lee, T. Lu, G.R. Stark, H.M. Shen, Z.G. Liu, J. Park, and G.M. Hur. 2013. PHF20 regulates NF-kappaB signalling by disrupting recruitment of PP2A to p65. Nature communications 4:2062.

Zhao, Y., Y. Lin, H. Zhang, A. Manas, W. Tang, Y. Zhang, D. Wu, A. Lin, and J. Xiang. 2015. Ubl4A is required for insulin-induced Akt plasma membrane translocation through promotion of Arp2/3-dependent actin branching. Proc Natl Acad Sci U S A 112:9644–9649.

Zhong, H., R.E. Voll, and S. Ghosh. 1998. Phosphorylation of NF-kappa B p65 by PKA stimulates transcriptional activity by promoting a novel bivalent interaction with the coactivator CBP/p300. Mol Cell 1:661–671.

